# Novel Insights into T-Cell Exhaustion and Cancer Biomarkers in PDAC Using ScRNA-Seq

**DOI:** 10.1101/2025.01.13.632574

**Authors:** Muhammad Usman Saleem, Hammad Ali Sajid, Muhammad Imran Shabbir, Alejandro Rivera Torres, Sunil Kumar Rai, Muhammad Waqar Arshad

## Abstract

One of the aggressive and lethal cancers, Pancreatic ductal adenocarcinoma (PDAC) is characterised by poor prognosis and resistance to conventional treatments. Moreover, tumor immune microenvironment (TIME) plays a crucial role in the progression and therapeutic resistance of PDAC. It is associated with T-cells exhaustion, leading to the progressive loss of T-cell functions with impaired ability to kill tumor cells. Therefore, this study employed single cell RNA sequencing (scRNA-seq) analysis identifying upregulated genes of T-cells; and of cancer cells in two groups (“cancer cells_vs_all-PDAC” and “cancer-PDAC_vs_all-normal”). Common and unique markers of cancer cells from both groups were identified. The Reactome pathways of cancer and T-cells were selected; while the genes implicated in those pathways were used to perform PPI analysis, revealing the hub-genes of cancer and T-cells. The gene expression validation of cancer and T-cells hub-genes was performed using GEPIA2 and TISCH2, while overall survival analysis of cancer cells hub-genes was performed using GEPIA2. Conclusively, this study unravelled 16 novel markers of cancer and T-cells providing the groundwork for future research into the immune landscape of PDAC, particularly T-cell exhaustion. However, further clinical studies are needed to validate these novel markers as potential therapeutic targets in PDAC patients.

**Graphical abstract:** 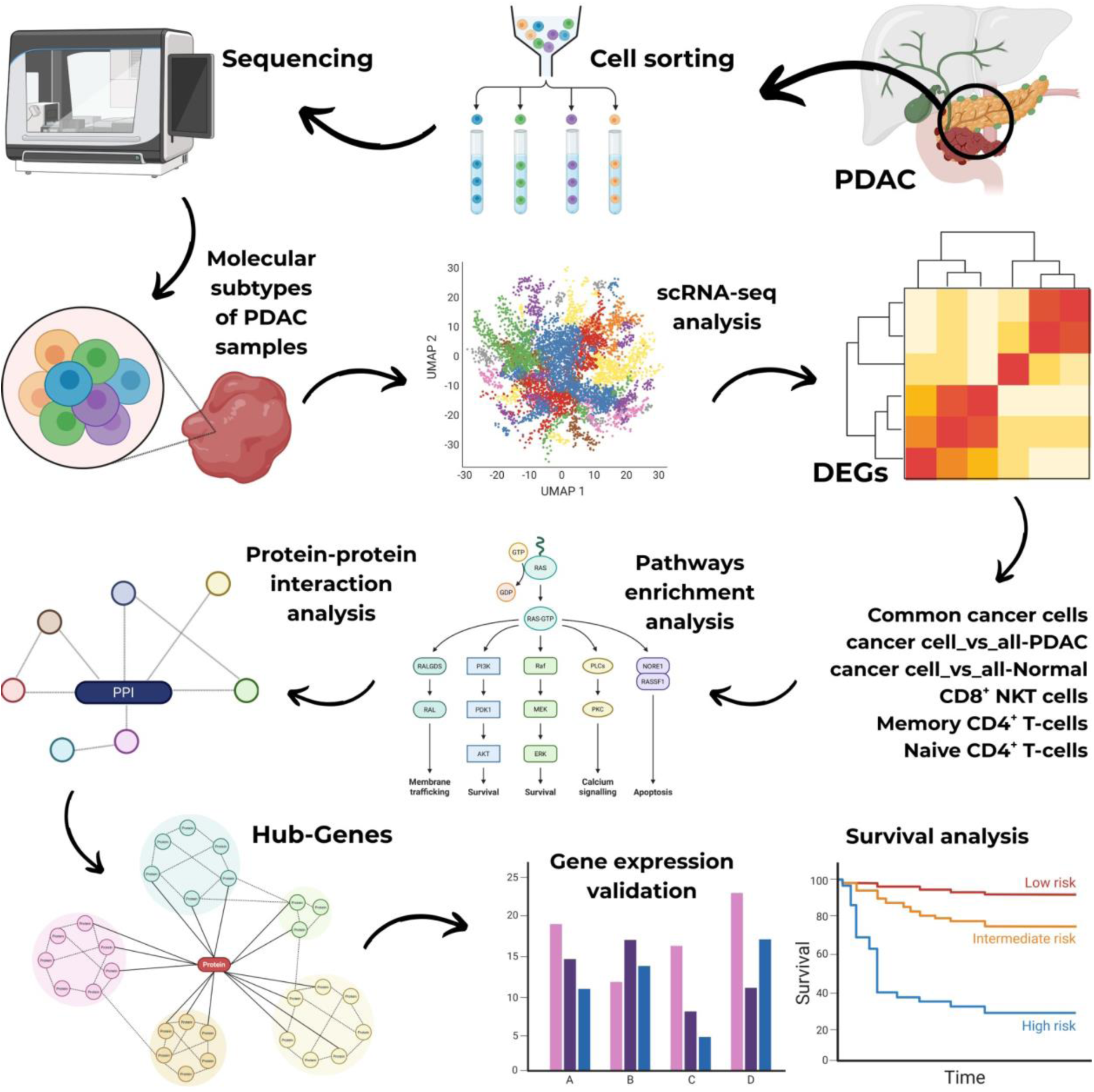

## 1. Introduction

Pancreatic tumors are classified as epithelial or non-epithelial based on their histogenesis, where pancreatic ductal adenocarcinoma (PDAC) is a most common malignant epithelial tumor of the pancreas accounting for more than 85% of all pancreatic malignancies [1]. The cancerous exocrine duct cells lining the pancreas leads to PDAC and is the deadliest of all adult abdominal tumors [2]. Moreover, PDAC has many subtypes, including classical and basal, squamoid-basaloid, treatment-enriched, and quasi-mesenchymal [3], [4].

Furthermore, PDAC is the third-leading cause of cancer death in the age groups 50–64 and 65–79; while there is only 10.8% 5-year overall survival rate for PDAC for both metastatic and resectable cases [5], [6]. Strikingly, 62,210 cases were reported in the United States in 2023 and 49,380 deaths in 2022, while PDAC accounts for 2% of all cancer cases and results in 5% of all cancer deaths in the United States, therefore underlining the crucial need for earlier detection [5], [7]. Notably, PDAC is estimated to become the second-leading cause of cancer deaths by 2030, surpassing breast cancer, as mortality rates are on the rise, increasing 1% annually [8].

Major risk factors of PDAC include smoking, obesity, diabetes, family history of pancreatic cancer, and inherited genetic mutations such as those in *BRCA1*, *BRCA2,* and *PALB2* [9], [10]. Subsequently, the common symptoms of PDAC patients include jaundice due to bile duct obstruction, unexplained weight loss, abdominal or back pain, and new-onset diabetes. These symptoms typically arise at advanced stages, contributing to the poor prognosis associated with PDAC [11].

Currently, different treatment regimens, including surgery, chemotherapy and radiation therapy are used to treat PDAC; however, these treatment options remain ineffective due to highly resistant PDAC [12]. Moreover, the gold standard to treat PDAC are polychemotherapy regimens, such as gemcitabine and abraxane or FOLFIRINOX [13]. Recently, advances in immunotherapy have led to a shift in treatment regimens by gaining insights into the intricate interactions between the immune system and cancer cells, while PDAC has been resistant to immunotherapy and has demonstrated marginal efficacy in terms of survival [14], [15].

Immunotherapy faces hurdles in PDAC treatment due to the immunosuppressive microenvironment that hinders the T-cell infiltration and activation, limiting the effectiveness of the combination therapy [16]. The complex tumor immune microenvironment (TIME) of PDAC modulates the infiltration of the immunosuppressive cells and the activity of immune regulatory molecules, leading to the dysfunctionality of the anti-tumor immune responses, including the exhaustion of the T-cells [17].

T-cell exhaustion is a dysfunctional state of the T-cells in chronic infections and cancer which results in cancer immune evasion, eventually leading to the poor prognosis in many cancers [18]. In PDAC, the T-cells get activated which later differentiate into memory-like cells, eventually leading to differentiated exhausted T-cells, which result from the persistent exposure of antigens on cells with low proliferative capacity, loss of their cytotoxic function, increased apoptosis, and elevated expression of multiple inhibitory receptors such as immune checkpoints, including PD-1, CTLA-4, TIM-3, LAG-3, or TIGIT [19], [20]. The mechanism of T-cell exhaustion induced within the PDAC TIME is illustrated in **Figure 1**.

**Figure 1.**
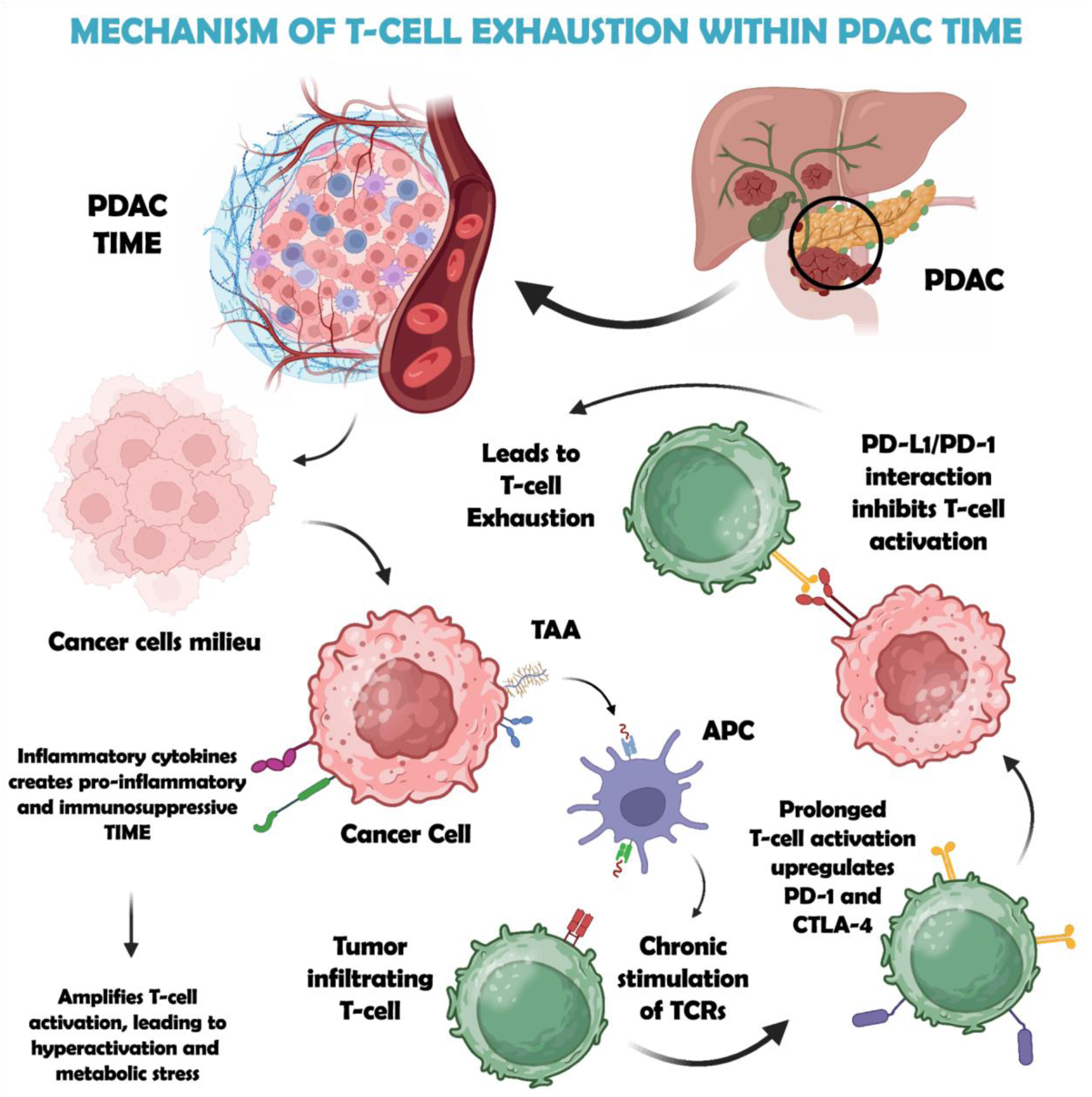
The mechanism of T-cell exhaustion induced within the PDAC TIME shows that in PDAC, cancer cells release tumor-associated antigens (TAAs) into the tumor immune microenvironment (TIME), which are presented by antigen-presenting cells (APCs) to T-cells, causing chronic TCR stimulation. This triggers T-cell hyperactivation, amplified by PDAC-secreted cytokines like IL-6 and TGF-β, leading to metabolic stress. Prolonged activation induces upregulation of inhibitory receptors (PD-1, CTLA-4) on T-cells, while PDAC cells overexpress PD-L1. The PD-L1/PD-1 interaction suppresses T-cell effector functions, reducing cytokine production (e.g., IFN-γ) and cytotoxicity, enabling tumor immune evasion.

Moreover, PDAC is characterised by tumor-infiltrating CD4^+^ and CD8^+^ T-cells; however as PDAC progresses, percentage of Tregs elevates within the CD4^+^ T-cell subset, while CD8^+^ T-cells shifts in a decreased composition [21], [22]. Although T-cells are observed within the TIME of PDAC, it is considered as a poorly immune responsive cancer, as T-cells exhibit lack of activation or an exhausted phenotype [23].

Furthermore, a recent scRNA-seq study in combination with multiplex immunohistochemistry imaging, evaluated the T-cell landscape in PDAC, demonstrating that infiltrated CD8+ T-cells exhibited senescent or an exhausted phenotype with high expression levels of TIGIT^+^ and CD39^+^ alongside PD-1^low/intermediate^ expression [24].

Another, scRNA-seq study revealed a reduction of ligand-receptor interactions after chemotherapy, especially between TIGIT on CD8^+^ T-cells and its receptors on cancer cells, identifying TIGIT as a key inhibitory checkpoint molecule, suggesting that chemotherapy may contribute to immunotherapy resistance by impacting the immune interactions within the PDAC TIME [25].

Therefore, this study aimed to unravel the immune landscape, specifically T-cell landscape to identify the novel exhaustive T-cell markers, driven by the markers of cancer cells. This study utilised computational methods such as scRNA-seq analysis, differential gene expression, protein-protein interaction analysis, hub-genes identification, and expression validations of the identified hub-genes to gain insights into the molecular and functional aspects of T-cell exhaustion and to identify novel markers of cancer and T-cells associated with T-cell exhaustion. However, further clinical research is needed to validate the findings in this study.

## 2. Materials and Methods

### 2.1. ScRNA-seq dataset retrieval

The preprocessed (mapped) scRNA dataset of PDAC with an accession id <u>GSE212966</u> was retrieved using the database, Gene Expression Omnibus (GEO) datasets (https://www.ncbi.nlm.nih.gov/gds) of National Center for Biotechnology Information (NCBI), a public repository that archives and distributes functional genomic data [26]. The dataset consisted of two conditions (PDAC and control), with six PDAC samples (T1-T6) and six control samples (N1-N6), comprising a total of 12 samples taken from untreated PDAC patients and sequenced using Illumina NovaSeq 6000 platform.

### 2.2. Data preprocessing and clustering

The “Seurat” package within R was utilised for the scRNA-seq analysis of the retrieved dataset, comprising 36601 genes and 57167 cells which were subjected to the quality control measures, including “nFeature_RNA > 200 & < 6000”, “nCount_RNA > 1000”, and mitochondrial reads < 10, filtering out the undesired and dead cells.

Moreover, the “LogNormalize” method was used for the normalization of the data, followed by the identification of 2000 highly variable features using the “variance stabilizing transformation (vst)” method. Furthermore, the data was scaled and linear dimensionality reduction was performed through principal component analysis (PCA). Subsequently, elbow plot was observed and first 15 PCs were utilised to identify the cell clusters using “FindNeighbors” and “FindClusters” functions at a resolution of 0.6 and utilising the Louvain algorithm.

### 2.3. Cell type annotation

The identified cell clusters were annotated using the “ScType” database (https://sctype.app/), a computational platform which provides automated cell-type identifications [27]. The “gene_sets_prepare” function was utilized to retrieve immune system specific gene sets from the ScType database, which prepared both positive and negative marker gene sets. Subsequently, “sctype_score” function was used to perform annotation based on the expression of prepared marker genes by computing the cell-type-specific scores. These scores were aggregated at the cluster level, allowing for the identification of the most likely cell type associated with each cluster. Additionally, clusters with scores lower than one-fourth of the number of cells in the respective cluster were labeled as “Unknown” to ensure annotation confidence.

### 2.4. Molecular subtypes classification of PDAC samples

Subsequently, a subset of cancer cells from PDAC condition was created to identify the molecular subtypes of all six tumor samples (T1-T6). This was performed to identify the heterogeneity of tumor samples, which can influence the tumor behaviour, treatment response and prognosis.

This was achieved by retrieving the marker genes of five molecular subtypes of PDAC, including classical/pancreatic progenitor, basal-like/squamous/quasi-mesenchymal, immunogenic, aberrantly differentiated endocrine exocrine (ADEX), and activated & normal stroma/stroma-rich, through extensive literature. The list of marker genes for all five molecular subtypes is mentioned in **Supplementary sheet 1**.

Furthermore, an R package, “UCell” (https://github.com/carmonalab/UCell) was utilized to evaluate the gene signatures (retrieved marker genes) in tumor samples (T1-T6). It applies the “Mann-Whitney U statistic” to compute the UCell signature scores which are robust to data size and heterogeneity. The UCell scoring was applied using the “AddModuleScore_UCell” function, and the enrichment scores for each gene signature were calculated independently for every cell in the cancer cells subset, predicting the molecular subtype of each tumor sample.

### 2.5. Gene expression profiling across conditions

Moreover, after the identification of cell types, DEGs of cancer cells, CD8+ NKT-like cells, memory CD4+ T cells, and naive CD4+ T cells were identified using the “FindMarkers” function and the “wilcox” method with a threshold of ±0.25.

The differential gene expression analysis of cancer cells was performed in two groups, including cancer cells compared to all other cell types (cancer cells_vs_all-PDAC) within the PDAC condition and cancer cells within PDAC condition compared to all other cell types in normal condition (cancer-PDAC_vs_all-normal). Moreover, the CD8+ NKT-like cells, memory CD4+ T cells, and naive CD4+ T cells within PDAC condition were compared to the same cells in normal condition (CD8+ NKT-like cells-PDAC_vs_CD8+ NKT-like cells-normal, memory CD4+ T cells-PDAC_vs_memory CD4+ T cells-normal, and naive CD4+ T cells-PDAC_vs_naive CD4+ T cells-normal).

The differential gene expression of the aforementioned combinations of cancer cells was performed to gain insights into the upregulated markers implicated in the heterogeneity and progression of cancer cells by exhausting the T-cells within the TIME. Furthermore, the differentially expressed genes of T-cells (CD8+ NKT-like cells, memory CD4+ T cells and naive CD4+ T cells) indicated the implication of upregulated markers in the loss of function of T-cells, which might be due to T-cell exhaustion, enabling cancer cells to grow and progress rapidly in TIME of PDAC.

Subsequently, common and unique upregulated markers of cancer cells from both groups were distinguished by utilising the Venn diagrams using the web-based tool, Venny 2.0 (https://bioinfogp.cnb.csic.es/tools/venny/index2.0.2.html).

### 2.6. Pathways enrichment analysis, protein-protein interaction (PPI) analysis and hub-genes identification

The identified upregulated genes of cancer cells, CD8+ NKT-like cells, memory CD4+ T cells, and naive CD4+ T cells were used for the pathways enrichment analysis using the GeneCodis4 (https://genecodis.genyo.es/), a web-based tool for the functional enrichment analysis allowing researchers to integrate different sources of annotations [28]. The Reactome pathways were selected and the genes implicated in the top-10 upregulated pathways were used for PPI analysis using STRING (https://string-db.org), a database which collects, scores and integrates all the publicly available sources of PPI information, and to complement these with the computational predictions [29].

Lastly, the top-10 enriched hub-genes were identified from the proteins interaction network through the “Degree” method by using the Cytoscape, an open-source software which integrates the biomolecular interaction networks with high-throughput expression data [30].

### 2.7. Aberrant genes expression validation and survival rates in PDAC patients

The gene expression validations and survival analysis of the identified hub genes of cancer cells was performed using GEPIA2 (http://gepia2.cancer-pku.cn/#index) web server, which is used for large-scale expression profiling and interactive analysis [31]. Moreover, the genes expression validation of CD8+ NKT-like cells, memory CD4+ T-cells, and naive CD4+ T-cells was performed using Tumor Immune Single Cell Hub 2 (TISCH2, http://tisch.comp-genomics.org/), a scRNA-seq data resource from human and mouse tumors, enabling extensive characterization of gene expression in TIME [32]. Furthermore, this was performed to shortlist the crucial hub genes implicated in the progression of PDAC.

## 3. Results

### 3.1. Data preprocessing and cell type annotation

The retrieved dataset consisting of six PDAC samples and six control samples were merged using the “merge” function in the “Seurat” package, resulting in a combined matrix. This matrix was then converted into a “Seurat Object”, which was further used for quality control measures retaining 38589 cells comprising 36601 genes. Subsequently, normalization, scaling, and highly variable features were identified. Among the 2000 highly variable features, the top-10 were *JCHAIN, IGKC, SPP1, EREG, APOC1, MZB1, IGHG1*, *TPSB2, CPA3* and *TPSAB1*. Moreover, the linear dimensionality reduction and cell clustering was performed based on gene expression profiles, resulting in a total of 26 cell clusters at a resolution of 0.6. The quality control, highly variable features, PCA plot, elbow plot, and t-SNE of cell clusters are shown in **Supplementary Figure S1(a-e)**.

Furthermore, the cell annotation resulted in 15 cell types, including Basophils, Cancer cells, CD8+ NKT-like cells, Endothelial, Macrophages, Memory CD4+ T cells, Myeloid Dendritic cells, Naive CD4+ T cells, Natural killer cells, Neutrophils, Plasma B cells, Plasmacytoid Dendritic cells, Platelets, Pre-B cells, and Progenitor cells. Interestingly, it was observed that the control condition exhibited trace amounts of cancer cells, indicating the spread of PDAC to control (healthy) regions of the pancreas, leading to metastasis.

Subsequently, it was observed that CD8+ NKT-like cells and memory CD4+ T cells exhibited a large and dense cluster in PDAC condition as compared to the control condition, suggesting the infiltration of T-cells at the tumor site. However, naive CD4+ T cells were observed to have a large and dense cluster of cells in control condition. Additionally, the cancer cells subset was created to identify the molecular subtypes of PDAC samples. The annotated cell types in control and PDAC conditions and the cancer cells subset are shown by using t-SNE method in **Figure 2 (a-b)**.

**Figure 2.**
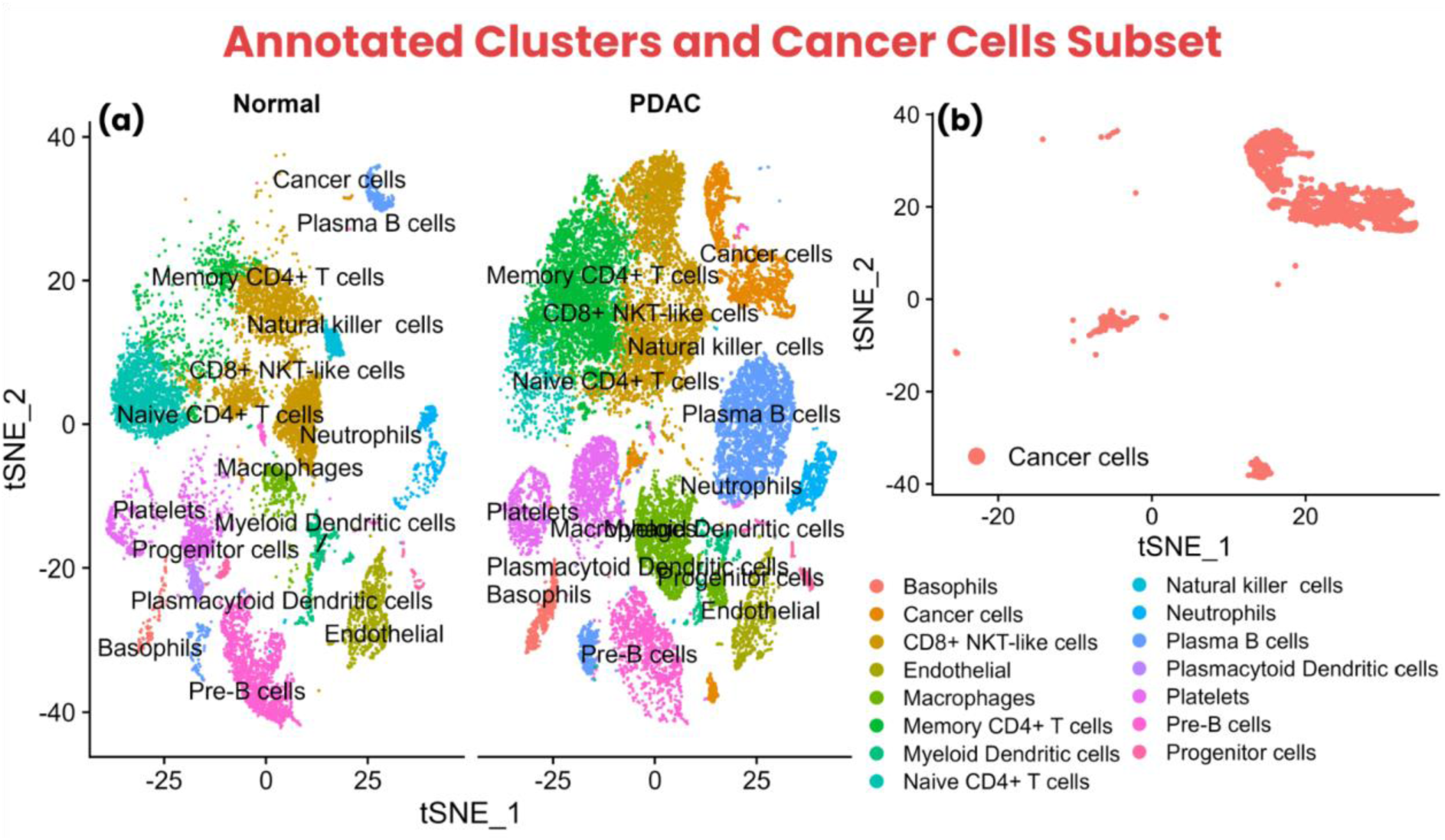
Cell type annotation clusters shown using t-SNE plot. **(a)** The annotated clusters within Normal and PDAC conditions showing different cell populations in both conditions, indicating complex influence of cancer cells in PDAC condition compared to Normal condition, **(b)** The cancer cells subset to identify molecular subtypes of PDAC tumor samples (T1-T6)

### 3.2. Molecular subtypes classification of PDAC samples

The identification of molecular subtypes indicated heterogeneous profiles of PDAC tumor samples, showing the disparity of tumor samples across multiple molecular subtypes. It was observed that the signature genes of “activated & normal stroma/stroma-rich” molecular subtype showed distributed cancer cells in trace amounts with moderate expression levels except for T5 and T6 samples, which showed low expression levels. However, the T3 and T4 samples showed moderate expression levels with moderately distributed cancer cells.

Interestingly, the signature genes of “classical/pancreatic progenitor”, “basal-like/squamous/quasi-mesenchymal”, “immunogenic”, and “ADEX” showed diverse amounts of cancer cells with varying expression levels in all six tumor samples (T1-T6), indicating heterogeneous samples.

The “immunogenic” and “ADEX” signature genes showed moderate cancer cells with low expression levels; however, the signature genes of “classical/pancreatic progenitor” and “basal-like/squamous/quasi-mesenchymal” showed highly-distributed dense cancer cells and high expression levels, with cancer cells being slightly more shifted towards “classical/pancreatic progenitor” molecular subtype in all six tumor samples as compared to “basal-like/squamous/quasi-mesenchymal”. The cancer cells distribution and expression levels of signature genes of respective molecular subtypes in individual tumor samples is shown in **Figure 3**.

**Figure 3.**
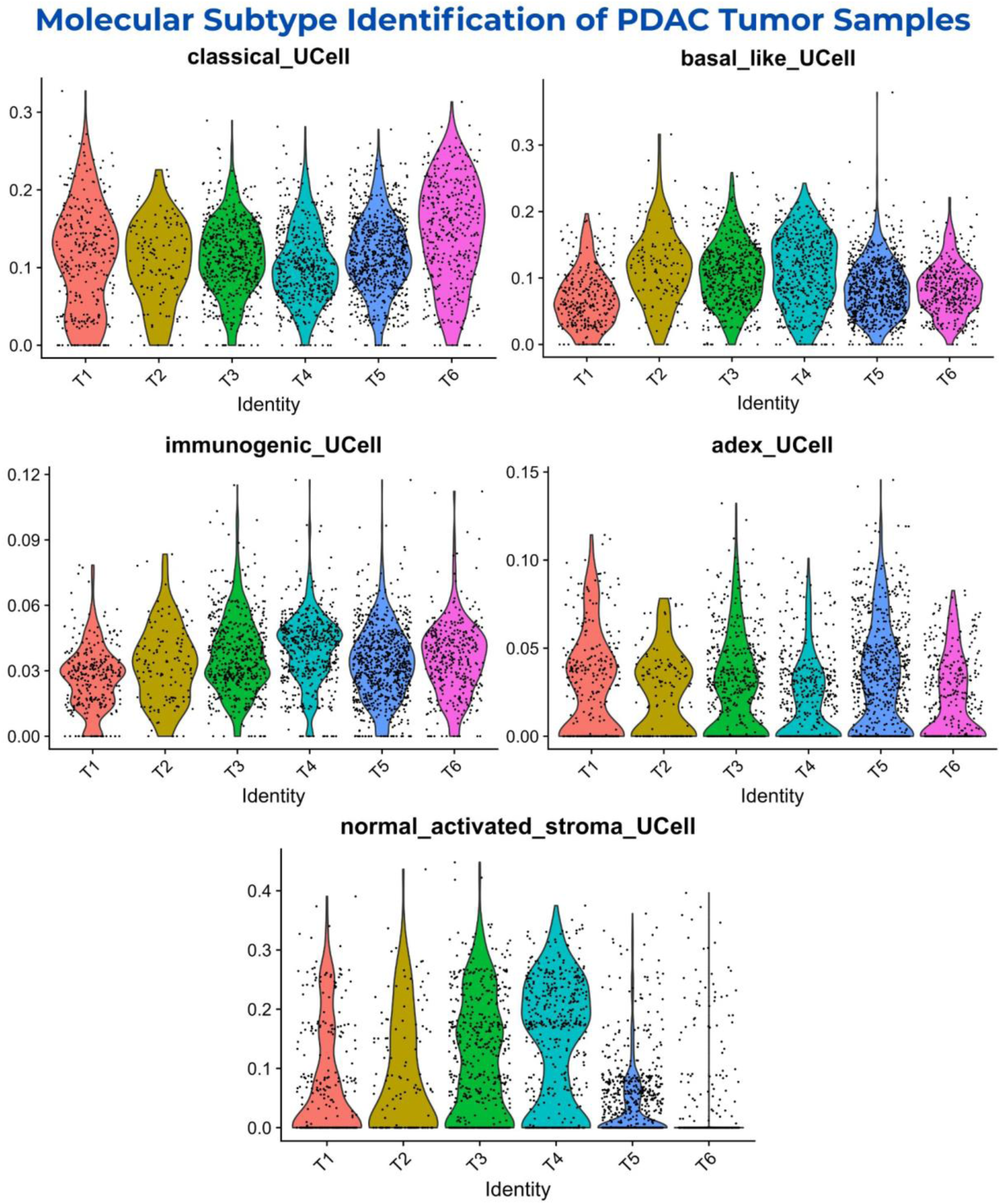
The identification of “classical/pancreatic progenitor”, “basal-like/squamous/ quasi-mesenchymal”, “immunogenic”, “ADEX”, and “activated & normal stroma/stroma-rich” molecular subtypes of PDAC tumor samples (T1-T6). The signature genes of “classical/pancreatic progenitor” and “basal-like/squamous/quasi-mesenchymal” showed expression levels ranging from 0 to 0.3, while “immunogenic” and “ADEX” ranged from 0 to 0.12 and from 0 to 0.15. Lastly, the signature genes of “activated & normal stroma/stroma-rich” showed expression levels ranging from 0 to 0.4 with trace amounts of cancer cells. This indicated a heterogeneity of all six tumor samples in the dataset

This suggests that all the tumor samples within the dataset showed heterogeneous nature with “classical/pancreatic progenitor” and “basal-like/squamous/quasi-mesenchymal” molecular subtypes being highly expressed in majority of the cancer cells, revealing aggressive nature of tumor samples with diverse genetic makeup. The expression levels of signature genes of respective molecular subtypes in cancer cells subset is shown in t-SNE plot in **Supplementary Figure S3**.

### 3.3. Aberrant markers of cancer cells, CD8+ NKT-like cells, memory CD4+ T cells and naive CD4+ T cells

The differential gene expression of the “cancer cells_vs_all-PDAC” group resulted in 8,509 upregulated and 4,642 downregulated genes, totalling to 13,151 dysregulated genes. Among these genes the top 10 upregulated included *AC007529.2, LINC02747, FGA, LGALS7, RASSF10, FER1L6, AC036176.3, NPSR1, SERPINB3,* and *AADAC* with p-values of 3.31E-47, 8.77E-49, 4.02E-75, 3.06E-49, 6.12E-60, 0, 6.53E-121, 0, 0, and 0; while logarithm of fold change (log2FC) of 8.60, 8.12, 7.89, 7.88, 7.82, 7.76, 7.75, 7.72, 7.71, and 7.63, respectively.

Moreover, the “cancer-PDAC_vs_all-normal” group exhibited 9,268 upregulated and 4,221 downregulated genes, totalling to 13,489 dysregulated genes. Among these genes the top 10 upregulated included *CEACAM5, MUC5B, SERPINB3, KRT6B, TFF2, AL121761.1, TFF1, CEACAM6, EPS8L3,* and *SPRR3* with p-values of 0 except for *SPRR3* showing a p-value of 1.58E-108; while these genes showed log2FC of 14.42, 14.30, 13.77, 13.47, 13.38, 13.11, 13.10, 13.07, 13.01, and 12.60, respectively. The expression levels of both groups of cancer cell genes are shown in **Supplementary Figure S4 (a-b)**.

Furthermore, CD8+ NKT-like cells exhibited 2,598 upregulated and 3,771 downregulated genes, totalling to 6,369 dysregulated genes. The top 10 upregulated genes included *COL11A1, MUC5AC, CEACAM5, FXYD3, MUC5B, MUC4, KLK6, TSPAN1, GPX2,* and *CTSE* with p-values of 1.78E-26, 5.76E-30, 9.61E-21, 3.25E-283, 1.35E-16, 2.17E-15, 1.03E-09, 1.65E-08, 1.90E-06, and 2.55E-28, while log2FC of 8.81, 8.80, 8.40, 8.23, 8.01, 7.94, 7.35, 7.34, 7.34, and 7.24 respectively.

The memory CD4+ T cells showed 2,820 upregulated and 3,500 downregulated genes, totalling to 6,320 dysregulated genes. The top 10 upregulated genes included *FXYD3, TFF1, MUC1, OLFM4, AGR2, LCN2, CTSE, CEACAM6, KRT19,* and *CLDN4* with p-values of 1.53E-18, 1.22E-14, 1.33E-11, 7.02E-08, 1.90E-05, 6.27E-05, 5.09E-03, 3.64E-03, 7.40E-13, and 2.02E-04; while log2FC of 8.42, 8.17, 7.51, 7.45, 6.97, 6.75, 6.51, 6.47, 6.37, and 5.39, respectively.

The naive CD4+ T cells resulted in 4,331 upregulated and 3,012 downregulated genes, totalling to 7,343 dysregulated genes. The top 10 upregulated genes included *KRT19, TFF1, LCN2, AGR2, TFF2, OLFM4, CLDN18, SPP1, C19orf33,* and *CEACAM6* with p-values of 1.14E-19, 4.28E-10, 7.87E-11, 4.89E-13, 1.45E-11, 3.76E-07, 1.45E-11, 2.12E-33, 7.87E-11, and 4.28E-10; while log2FC of 8.30, 7.95, 7.81, 7.75, 7.72, 7.61, 7.49, 7.49, 7.46, and 7.38, respectively. The expression levels of T-cells top 10 upregulated markers are shown in **Supplementary Figure S4 (c-e)**.

It was observed that the “cancer-PDAC_vs_all-normal” group exhibited the most and high intensity of dysregulation compared to the “cancer cells_vs_all-PDAC” group, indicating that markers in this group might be implicated in the rapid progression of cancer cells in TIME and may be highly implicated in signalling, leading to T-cell exhaustion. Nevertheless, it was observed that among T-cells, naive CD4+ T cells showed the highest number of dysregulated genes, indicating high influence of cancer cells on naive CD4+ T cells.

Subsequently, the upregulated markers of cancer cells showed 7,882 common genes among both groups, while 1,386 unique to “cancer-PDAC_vs_all-normal”; and 627 unique to “cancer cells_vs_all-PDAC” group. Henceforth, all these genes were used separately for further analysis to identify unique and common markers in cancer cells while comparing different conditions. The venn diagram of “cancer-PDAC_vs_all-normal” and “cancer cells_vs_all-PDAC” groups is shown in **Supplementary Figure S4 (f)**.

### 3.4. Pathways dysregulation leading to T-cell exhaustion and PDAC progression

The upregulated markers were used to identify the enriched Reactome pathways. The common markers of cancer cells included Metabolism, Metabolism of lipids, Asparagine N-linked glycosylation, Post-translational protein modification, Membrane Trafficking, RHO GTPase cycle, Vesicle-mediated transport, Regulation of cholesterol biosynthesis by SREBP (SREBF), ER to Golgi Anterograde Transport, and RHOB GTPase cycle to be enriched top-10 pathways.

While the unique markers to “cancer cells_vs_all-PDAC” group included Gene expression (Transcription), Generic Transcription Pathway, RNA Polymerase II Transcription, Metabolism of RNA, DNA Repair, Antiviral mechanism by IFN-stimulated genes, SMAC (DIABLO) binds to IAPs, SMAC (DIABLO)-mediated dissociation of IAP complexes, Abasic sugar-phosphate removal via the single-nucleotide replacement pathway, and SMAC, XIAP-regulated apoptotic response to be top-10 enriched pathways.

Lastly, the unique markers to “cancer-PDAC_vs_all-normal” group included Extracellular matrix organization, Collagen formation, Assembly of collagen fibrils and other multimeric structures, Collagen degradation, Collagen biosynthesis and modifying enzymes, Degradation of the extracellular matrix, ECM proteoglycans, Collagen chain trimerization, MET activates PTK2 signaling, and Integrin cell surface interactions to be enriched top-10 pathways. The Reactome pathways of cancer cells markers are shown in **Supplementary Figure S2 (a-c)**.

Moreover, the CD8+ NKT-like cells exhibited Extracellular matrix organization, Non-integrin membrane-ECM interactions, Assembly of collagen fibrils and other multimeric structures, Collagen formation, Signaling by Receptor Tyrosine Kinases, ECM proteoglycans, MET activates PTK2 signaling, Immune System, MET promotes cell motility, and Syndecan interactions top-10 upregulated pathways.

Furthermore, the top-10 upregulated pathways of memory CD4+ T cells included Signaling by Receptor Tyrosine Kinases, Neutrophil degranulation, Immune System, Syndecan interactions, Assembly of collagen fibrils and other multimeric structures, Potential therapeutics for SARS, Collagen degradation, Extracellular matrix organization, Disease, and TP53 Regulates Transcription of Genes Involved in G2 Cell Cycle Arrest.

Lastly, the top-10 enriched pathways of naive CD4+ T cells exhibited Gene expression (Transcription), Metabolism of RNA, Post-translational protein modification, Immune System, Processing of Capped Intron-Containing Pre-mRNA, RNA Polymerase II Transcription, Cytokine Signaling in Immune system, Vesicle-mediated transport, Membrane Trafficking, and mRNA Splicing. The upregulated Reactome pathways of T-cells markers are shown in **Supplementary Figure S2 (d-f)**.

Notably, it indicated that all the cancer cells pathways were uniquely implicated in cancer cells to evade the immune system and to promote its growth and progression in TIME by exhausting the T-cells. Moreover, it was observed that “Immune system” pathway was common among all T-cells, while “Extracellular matrix organization”, “Assembly of collagen fibrils and other multimeric structures”, “Signaling by Receptor Tyrosine Kinases”, and “Syndecan interactions” pathways were found to be common among CD8+ NKT-like cells and memory CD4+ T cells.

This suggested that cancer cells were sending similar signals to upregulate similar pathways in CD8+ NKT-like cells and memory CD4+ T cells. However, naive CD4+ T-cells did not exhibit common pathways with other T-cells except for the “Immune system”, suggesting the exhaustion of all three T-cells in this study, eventually leading to suppression of the immune system. This also indicates the heterogeneity that cancer cells dysregulated the immune cells (T-cells) through specific signals to specific T-cells, which creates complex interactions within the TIME of PDAC.

### 3.5. Key candidate proteins associated with T-cell exhaustion within PDAC TIME

The genes from top-10 significant pathways of each cell type were subjected to PPI analysis resulting in different networks of proteins interacting with each other. The upregulated 1,960 common pathway genes of cancer cells resulted in a network of 1,949 proteins interacting with each other. Moreover, 124 upregulated genes of the “cancer cells_vs_all-PDAC” group exhibited a network of 91 proteins, while 69 upregulated genes of the “cancer-PDAC_vs_all-normal” group resulted in a network of 69 proteins interacting with each other. The PPI networks of cancer cell genes are shown in **Supplementary Figure S5 (a-c)**.

Furthermore, 520 upregulated genes of CD8+ NKT-like cells exhibited a network of 513 proteins, while 657 enriched genes of memory CD4+ T cells showed a network of 639 proteins, and 1,454 upregulated genes of naive CD4+ T cells showed a network of 1,445 proteins. The protein networks of CD8+ NKT-like cells, memory CD4+ T cells, and naive CD4+ T cells are shown in **Supplementary Figure S5 (d-f)**.

Subsequently, these protein networks of each cell type were used to identify the most interacting genes (hub-genes) using the “Degree” method. The protein network of common cancer cell genes showed top-10 hub genes, including *GAPDH, AKT1, EGFR, CS, RHOA, TPI1, SDHA, TFRC, FASN,* and *HIF1A*; showing 364, 284, 242, 212, 207, 200, 189, 189, 186, and 179 interactions in the network, respectively.

The protein network of unique cancer cell genes of “cancer cells_vs_all-PDAC” group exhibited top 10 hub genes, including *H4C6* (*HIST1H4A* or *HIST1H4C*)*, MYC, H3C12* (*HIST1H3A* or *HIST1H3D*)*, DDX21, USP7, RFC4, APEX1, CDK9, H2BC9* (*HIST1H2BH*), and *NOP2*. Among these proteins, *RFC4, APEX1, CDK9, HIST1H2BH, and NOP2* showed 12 interactions, while *DDX21* and *USP7* showed 16 interactions; lastly, *H4C6, MYC* and *H3C12* exhibited 29, 28, and 18 interactions, respectively.

The protein network of unique cancer cell genes of “cancer-PDAC_vs_all-normal” group showed top 10 hub genes, including *FN1, COL1A1, COL1A2, COL3A1, COL5A2, COL6A1, COL5A1, BGN, COL6A2,* and *FBN1*, showing 62, 59, 56, 55, 51, 50, 50, 48, 46, and 45 interactions within the protein network, respectively. The hub-genes of cancer cells are shown in **Figure 4 (a-c)**. The common and unique hub genes of cancer cells from both groups are mentioned in **Supplementary Table S1**.

**Figure 4.**
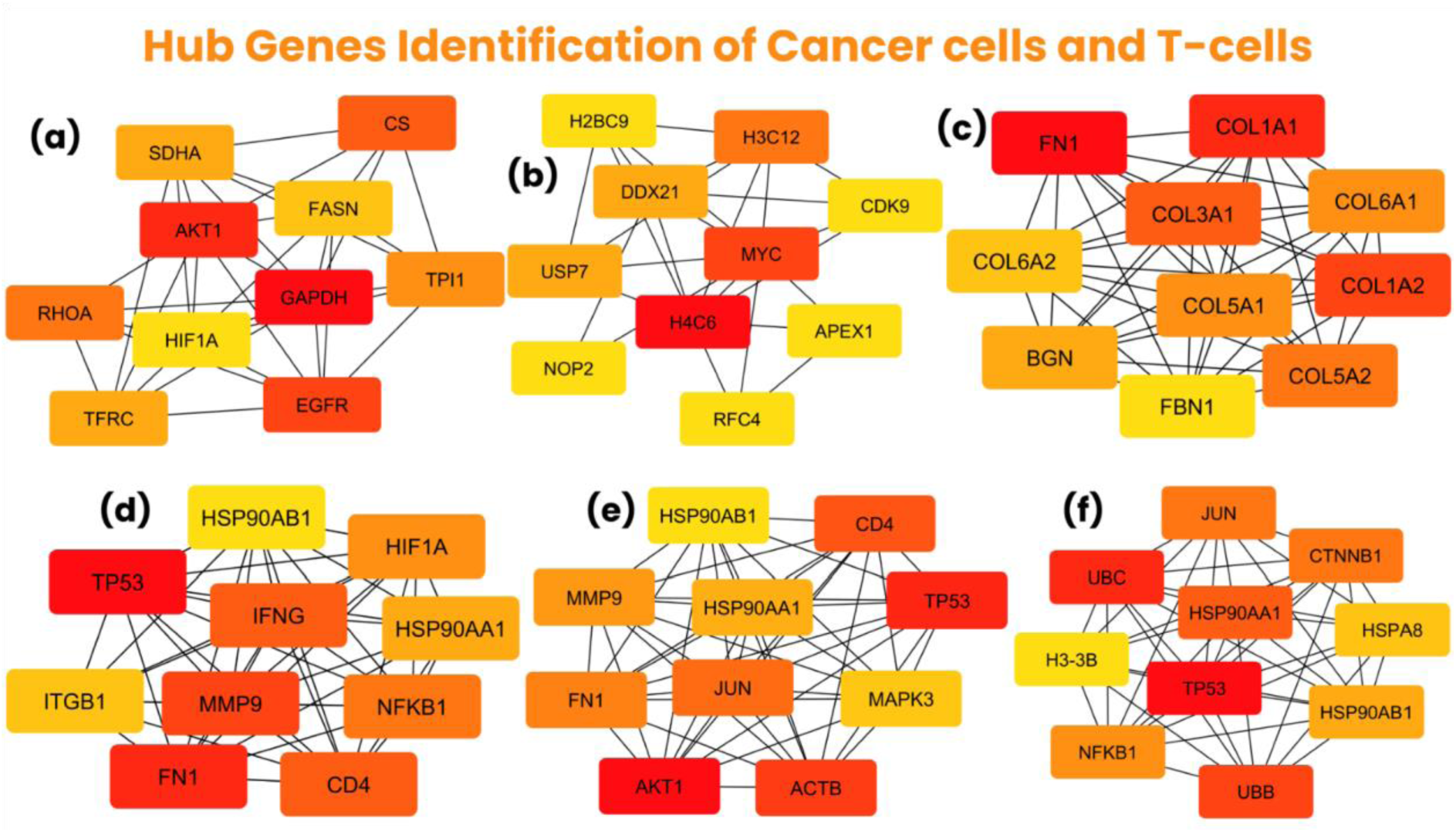
Hub genes identification of cancer cells and T-cells using Cytoscape. **(a)** The top 10 hub genes of “common cancer cells” group with *GAPDH* exhibiting the most interactions, **(b)** The top 10 hub genes of “cancer cells_vs_all-PDAC” group with *H4C6* showing the highest number of interactions, **(c)** The top 10 hub genes of “cancer-PDAC_vs_all-normal” group with *FN1* showing the most interactions, **(d)** The top 10 hub genes of CD8+ NKT-like cells with *TP53* as the most interacting hub gene, **(e)** The top 10 hub genes of memory CD4+ T cells with *AKT1* exhibiting most interactions, **(f)** The top 10 hub genes of naive CD4+ T cells with *TP53* as the most interacting hub gene

Moreover, the protein network of CD8+ NKT-like cells exhibited *TP53, FN1, MMP9, CD4, IFNG, NFKB1, HIF1A, HSP90AA1, ITGB1,* and *HSP90AB1* as the top 10 hub genes, showing 176, 170, 149, 147, 147, 142, 129, 126, 119, and 116 interactions, respectively. Furthermore, *AKT1, TP53, ACTB, CD4, JUN, FN1, MMP9, HSP90AA1, MAPK3,* and *HSP90AB1* were the top 10 hub genes in memory CD4+ T cells, exhibiting 200, 192, 180, 148, 146, 142, 126, 123, 121, and 113 interactions. Lastly, the protein network of naive CD4+ T cells showed *TP53, UBC, UBB, HSP90AA1, JUN, CTNNB1, NFKB1, HSP90AB1, HSPA8,* and *H3-3B* (*H3F3B*) top 10 hub genes, showing 452, 298, 279, 270, 260, 260, 251, 238, 229, and 212 interactions, respectively. The hub-genes of T-cells are shown in **Figure 4 (d-f)**. The hub genes of all three T-cells are mentioned in **Supplementary Table S2**.

Notably, the sub-groups of cancer cells (common genes, cancer cells_vs_all-PDAC, and cancer-PDAC_vs_all-normal) indicated the unique functional implication of respective hub-genes within their specific group. This suggests the complexity and heterogeneity of cancers within the TIME, as multiple different genes are implicated in the progression of PDAC cancer by several factors, including T-cell exhaustion.

Subsequently, it was observed that *TP53, HSP90AA1,* and *HSP90AB1* were common among all three T-cells; *FN1, MMP9,* and *CD4* were common in CD8+ NKT-like cells and memory CD4+ T cells; *NFKB1* was common in CD8+ NKT-like cells and naive CD4+ T cells, while *JUN* was common in memory CD4+ T cells and naive CD4+ T cells. This indicated that multiple hub-genes of T-cells were common among each other, suggesting that markers of cancer cells might be implicated in upregulating common markers of different T-cells through various pathways which are essential for these T-cells. The expression of hub genes of cancer cells and T-cells in heatmap is shown in **Figure 5**.

**Figure 5.**
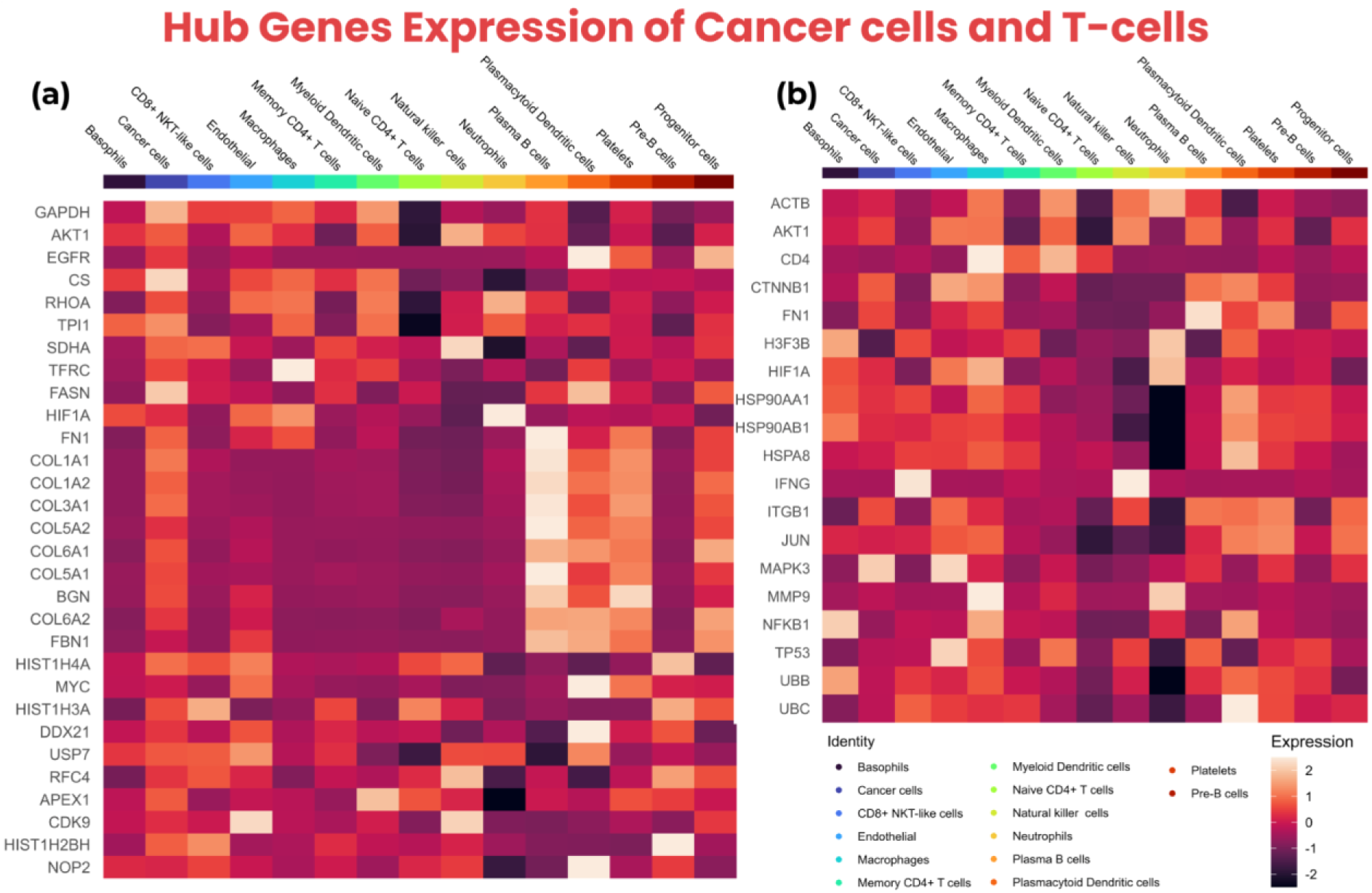
The hub genes expression of cancer cells and T-cells shown using heatmap. **(a)** The heatmap of top 10 hub genes from all three groups (common markers of cancer cells, cancer cells_vs_all-PDAC, and cancer-PDAC_vs_all-normal) showing high expression across multiple cell types, **(b)** The heatmap of top 10 hub genes of T-cells showing expression across multiple cell types including CD8+ NKT-like cells, memory CD4+ T cells, and naive CD4+ T cells

Additionally, the unique markers of T-cells indicated that cancer cells also influence those markers of T-cells which are unique to respective T-cell types, to evade the immune response through T-cell exhaustion. The expression of cancer cells and T-cells hub-genes across different cell types exhibited in dot plots, violin plots and box plots are shown in **Supplementary Figures S6-S8**.

### 3.6. Expression profiling and survival correlation of key hub-genes

The GEPIA2 was used for the expression validation and survival analysis of cancer cell hub genes as it contains TCGA bulk-RNA datasets, while TISCH2 was utilised for the expression validation of T-cells and cancer cells hub genes as TISCH2 contains the scRNA-seq datasets. Moreover, the GEPIA2 showed all cancer cells hub-genes to be highly upregulated in the tumor compared to the control samples. The expression values of all the hub genes of cancer cells in tumor and control samples are mentioned in **Supplementary Table S3**.

Furthermore, the survival analysis of the cancer cells hub-genes indicated that *GAPDH, AKT1, CS, RHOA, TPI1, SDHA, FASN, HIF1A, FN1, COL1A1, COL1A2, COL3A1, COL5A2, COL5A1, BGN, COL6A2, FBN1, USP7, CDK9, H2BC9,* and *NOP2* exhibited insignificant p-values, while *EGFR, TFRC, COL6A1, MYC, DDX21, RFC4,* and *APEX1* exhibited significant p-values including, 0.03, 0.04, 0.04, 0.01, 0.01, 0.04, and 0.04, respectively. Lastly, GEPIA2 showed no results for the overall survival analysis of *H4C6* and *H3C12*.

Subsequently, the significant hub-genes, including *EGFR* showed that high expression exhibited ∼18% of patients survival for more than 60 months while low expression showed ∼20% of patients survived for approximately more than 90 months. Moreover, the high expression of *TFRC* showed survival of ∼18% patients for more than 70 months, while low expression showed the survival of ∼18% patients for more than 90 months. Furthermore, the high expression of *COL6A1* indicated that none of the patients survived more than 70 months, while ∼37% of the low expression patients survived for more than 90 months.

The high expression of the MYC showed ∼11% of patients survived for more than 70 months, while ∼29% of low expression patients survived for more than 90 months. The high expression of the *DDX21* showed that none of the patients survived more than 75 months, while ∼38% of the low expression patients survived more than 90 months. The high expression of *RFC4* showed ∼13% of patients survived for ∼80 months, while ∼32% of patients survived for more than 90 months. Lastly, the high expression of *APEX1* showed that none of the patients survived for more than 75 months, while ∼35% of patients survived for more than 90 months. The overall survival plots of all the cancer cells hub-genes are shown in **Supplementary Figure S9**.

Additionally, the TISCH2 indicated that all the hub-genes of cancer cells were upregulated in malignant cells, especially *GAPDH, RHOA, TPI1, H4C6, DDX21,* and *APEX1* showed considerable high expression levels in malignant cells. Moreover, the hub-genes of T-cells indicated the upregulation of all hub-genes, especially *ACTB, H3F3B, HSP90AA1, HSP90AB1, HSPA8, UBB,* and *UBC* exhibited high levels of expressions in CD8Tex cells. The feature plots of cancer cells and T-cells hub genes from TISCH2 are shown in **Supplementary Figures S10 and S11**. The violin plots of cancer cells and T-cells hub-genes are shown in **Supplementary Figure S12**.

This indicated that the hub-genes of cancer cells and T-cells showed high expressions in datasets from GEPIA2 and TISCH2. However, the insignificant p-values in overall survival rate of the cancer cell hub-genes further need corroborations as these genes are well-reported to be expressed in PDAC patients leading to poor prognosis.

## 4. Discussion

There is a high rate of recurrence and metastasis in PDAC patients after surgery, lacking efficient chemotherapy, radiotherapy, and immunotherapy [33]. T-cell exhaustion, which is induced by the complex immunosuppressive TIME, may contribute to diminished responses of immunotherapy in PDAC patients [34]. Therefore, this study aimed to identify the upregulated markers of cancer cells and T-cells implicated in the PDAC progression through T-cell exhaustion by utilising scRNA-seq analysis.

Subsequently, this study revealed heterogeneous tumor samples, mainly classified to “classical/pancreatic progenitor” and “basal-like/squamous/quasi-mesenchymal” molecular subtypes, which provided significant insights into the underlying biological diversity of PDAC. Moreover, this study revealed enriched pathways of cancer cells implicated in crucial biological processes for maintaining cellular homeostasis, including lipid and protein metabolism, post-translational modifications, and membrane trafficking [35], [36], [37]. Moreover, pathways such as Regulation of gene expression, RNA metabolism, and DNA repair are essential for efficient cell function and survival [38], [39], [40]. This suggests that the cancer cells pathways collectively play a vital role in maintaining cellular integrity, communication, and responses to external signals, leading to cancer cells progression by evading the immune system.

Furthermore, the enriched pathways of CD8+ NKT-like cells, memory CD4+ T-cells, and naive CD4+ T-cells were implicated in gene regulation, cellular communication, and immune responses. The pathways, including Receptor tyrosine kinases and MET signaling pathways impact immune responses and cell movement by regulating cell motility, growth and survival [41], [42]. Additionally, the immune related pathways such as neutrophil degranulation and cytokine signaling were highly expressed in T-cells [43], [44]. This indicated that the high expression of immune related pathways might be leading to T-cell exhaustion, rendering them non-functional against cancer cells.

Notably, all three sub-groups (common, cancer cells_vs_all-PDAC, and cancer-PDAC_vs_all-normal) of cancer cells exhibiting hub-genes, indicated the heterogeneity and complexity of cancer cells within the TIME. Moreover, all the hub-genes, including *FN1, COL1A1, COL1A2, COL3A1, COL5A2, COL6A1, COL5A1, BGN, COL6A2,* and *FBN1* of “cancer-PDAC_vs_all-normal” group are reported to be implicated in PDAC patients leading to poor prognosis [45], [46], [47], [48], [49], [50], [51].

Similarly, the common hub-genes such as *GAPDH, AKT1, EGFR, RHOA, TPI1, SDHA, TFRC, FASN,* and *HIF1A* are reported to be implicated in oncogenic pathways leading to cancer cells proliferation and progression, playing crucial role in PDAC [52], [53], [54], [55], [56], [57], [58], [59], [60], while *CS* is not specifically reported in PDAC, resulting in a novel PDAC cancer cell marker.

Furthermore, the *MYC, DDX21, USP7, RFC4, APEX1, CDK9,* and *NOP2* of “cancer cells_vs_all-PDAC” group play critical role in the metabolic reprogramming of the PDAC, contributing to poor prognosis of PDAC [61], [62], [63], [64], [65], [66], [67]. However, *H4C6, H3C12*, and *H2BC9* are not reported in cancer cells of PDAC patients and these novel markers might be implicated in cell proliferation and survival of cancer cells by sending signals, leading to exhaustion of T-cells.

Additionally, among the common hub-genes of all three or any two T-cell subtypes (CD8+ NKT-like cells, memory CD4+ T-cells, and naive CD4+ T-cells), *CD4* is reported to be activated in T-effs and strongly promote EMT-associated alterations in H6c7 cells and increased invasive behaviour [68]. NFKB1 within the NFKB signaling is influenced in cytotoxic T-cells in the blood of patients with PDAC [69]. *HSP90AA1* has been reported to be expressed in a new type of T-cell (HSP T or Thsp) within the PDAC environment [70], while *JUN* is reported to be implicated in the T-cell exhaustion mechanism within the PDAC environment [71].

However, *TP53, MMP9*, *FN1,* and *HSP90AB1* were observed to be novel markers as these markers are not specifically reported in CD8+ NKT-like cells, memory CD4+ T-cells, naive CD4+ T-cells or any related T-cell within PDAC patients, indicating the involvement of these novel markers in T-cell exhaustion and aberrant functional activity within PDAC TIME, eventually leading to PDAC cells proliferation and survival.

Moreover, the unique hub-genes of CD8+ NKT-like cells showed that *IFNG* and *HIF1A* are reported to be implicated in T-cells [72], [73], while *ITGB1* turned out to be a novel marker of CD8+ NKT-like cells. Furthermore, the unique hub-genes of memory CD4+ T-cells such as *AKT1, ACTB*, and *MAPK3* were observed to be novel markers as these are not reported specifically in T-cells of PDAC patients. Lastly, the unique hub-genes of naive CD4+ T-cells only *HSPA8* is reported to be expressed in CD8+ T-cells in prostate cancer [74], while *UBC, UBB, CTNNB1*, and *H3-3B* are not reported, rendering them novel markers for naive CD4+ T-cells. This suggests that the aforementioned novel markers are potentially implicated in exhaustion of T-cells within TIME, eventually leading to cancer cells growth, proliferation, and survival by evading the immune responses in PDAC patients.

Subsequently, the expression and survival analysis of the cancer cells and T-cells hub-genes exhibited majority of the cancer cells overall survival to be insignificant (p-value > 0.05); however, aforementioned studies support the aberrant expression of these markers to be implicated in poor prognosis in PDAC patients. Additionally, further corroboration is needed, especially for novel markers of cancer cells and T-cells within PDAC patients for diagnostic, prognostic, and therapeutic interventions.

Recent scRNA-seq studies on T-cells within PDAC patients have reported T-cell marker genes in CD8+ Tcm, CD4+ Tem, and γδT cells; and reported that the *CCL5/SDC1* receptor-ligand interactions in tumor infiltrating T-cells could promote tumor cells migration [75], [76]. However, this study reveals novel biomarkers of cancer cells, CD8+ NKT-like cells, memory CD4+ T-cells, and naive CD4+ T-cells implicated in T-cell exhaustion, rendering the growth, proliferation and progression of cancer cells.

In summary, the aforementioned novel biomarkers of heterogeneous cancer cells and T-cells might be involved in the complex TIME, leading to the poor prognosis of PDAC. These findings underscore the complexity and heterogeneity of the TIME and its influence on immune evasion and cancer cell survival. While existing studies support the role of some identified markers in poor prognosis, further clinical validation is required for PDAC management.

## 5. Conclusion

This study revealed novel biomarkers of heterogeneous cancer cells (*CS, H4C6, H3C12,* and *H2BC9*) and T-cells (*TP53, FN1, MMP9, ITGB1, HSP90AB1, AKT1, ACTB, MAPK3, UBC, UBB, CTNNB1,* and *H3-3B*), including CD8+ NKT-like cells, memory CD4+ T-cells, and naive CD4+ T-cells, which might be the key candidate biomarkers implicated in cancer cells progression through T-cell exhaustion. Further clinical investigations are needed to evaluate these novel biomarkers as potential therapeutic options in PDAC patients.

## Supporting information

Supplementary Sheet 01

