## Supplementary Document 01 for "Novel Insights into T-Cell Exhaustion and Cancer Biomarkers in PDAC Using ScRNA-Seq"

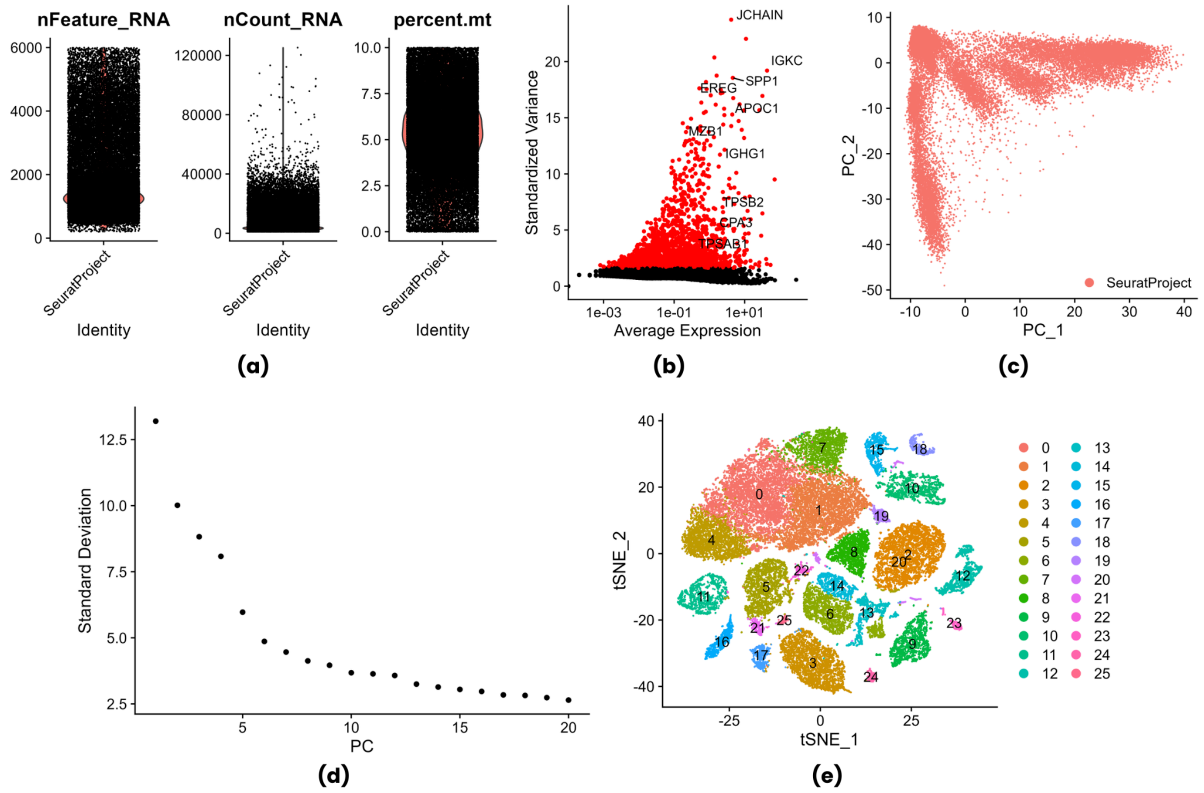


**Figure S1.** ScRNA-seq dataset preprocessing steps showing **(a)** filtered cells based on number of features, RNA counts, and mitochondrial reads; **(b)** 2000 highly variable features with labelled top-10 features; **(c)** PCA plot; **(d)** Elbow plot showing the variable principle components; and **(e)** t-SNE map of the clusters identified at resolution of 0.6

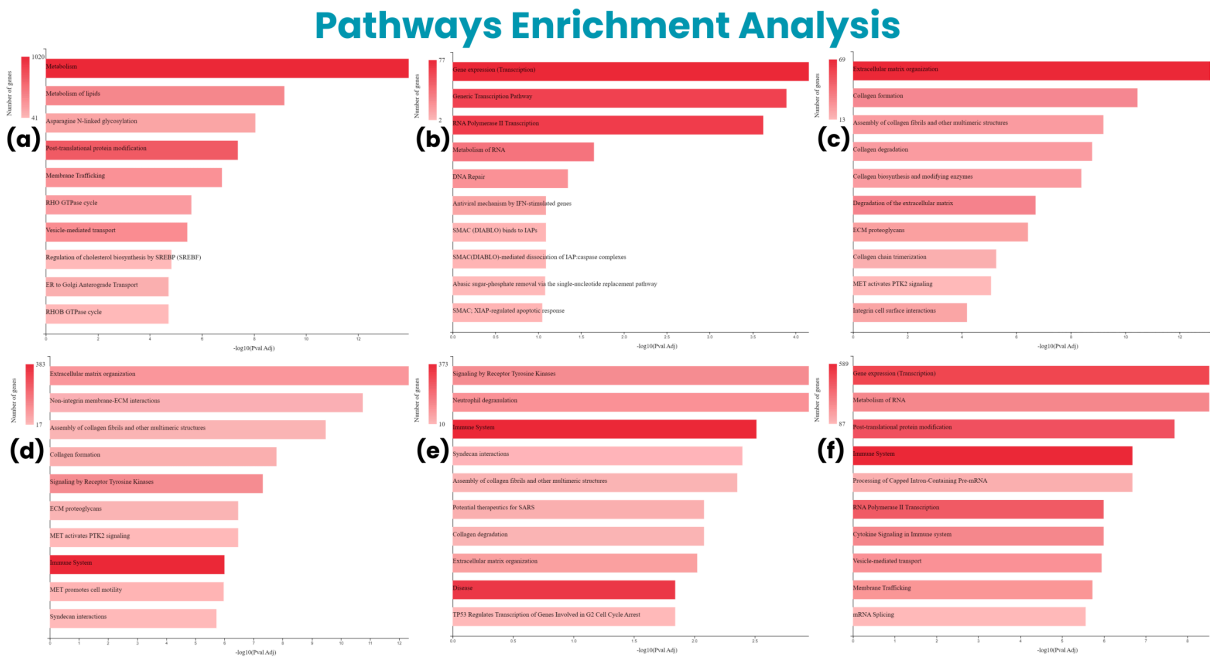


**Figure S2.** The enriched pathways implicated in cancer cells and T-cells. **(a-c)** Tp[ 10 upregulated Reactome pathways of cancer cells genes, **(d-f)** Top 10 upregulated Reactome pathways of CD8+ NKT-like cells, memory CD4+ T-cells, and naive CD4+ T-cells

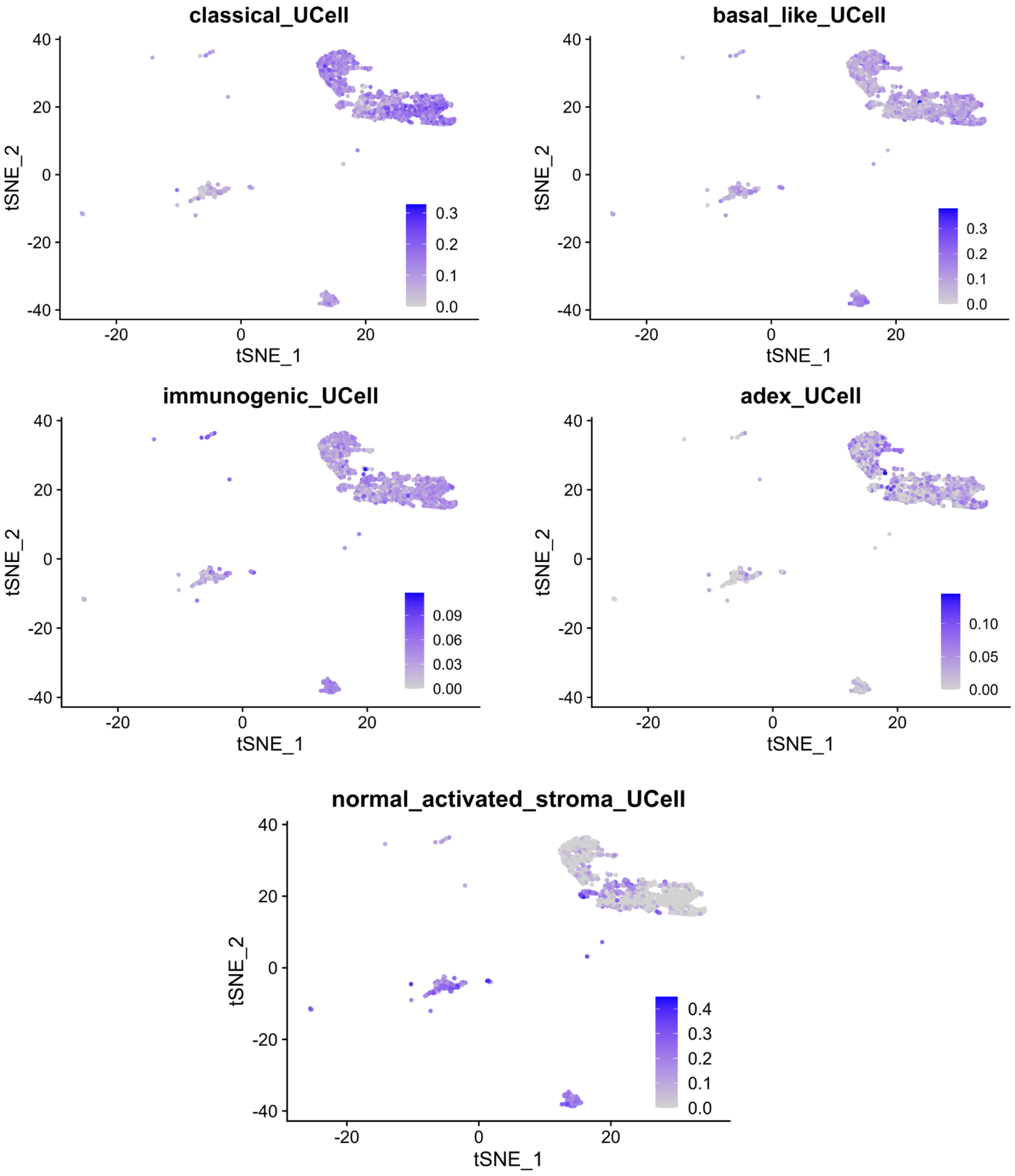


**Figure S3.** The t-SNE plots showing molecular subtypes identification in cancer cells subset from all tumor samples (T1-T6). The "classical/pancreatic progenitor" and "basal-like/squamous/quasi-mesenchymal" showed expression levels up to 0.3 with slightly more cells expressing signature genes of "classical/pancreatic progenitor". The "immunogenic" and "ADEX" showed expression levels up to 0.09 and 0.10, respectively, with moderately expressing cancer cells. Lastly, the "activated & normal stroma/stroma-rich" showed high expression level in trace amounts of cancer cells

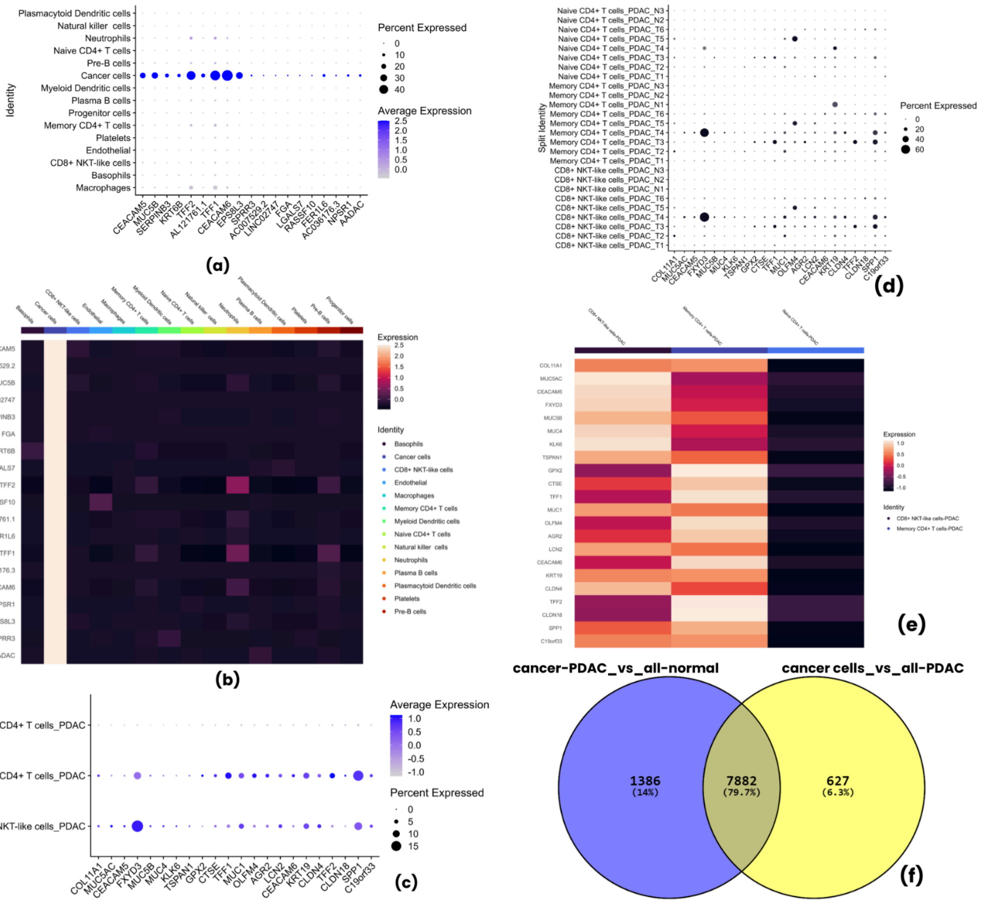

**Figure S4.** Expression plots of top-10 cancer and T-cells genes showing **(a-b)** dot plot and heatmap of the top-10 cancer cell genes, respectively; **(c)** dot plot of top-10 T-cells genes; **(d)** dot plot of top-10 T-cells genes in individual PDAC and normal samples, indicating slight expression of genes within Naive CD4+ T-cells, leading to overall low expression, **(e)** as also seen in heatmap; **(f)** Venn diagram of cancer cells genes of "cancer-PDAC_vs_all-normal" and "cancer cells_vs_all-PDAC" groups, indicating the common and unique genes among both groups

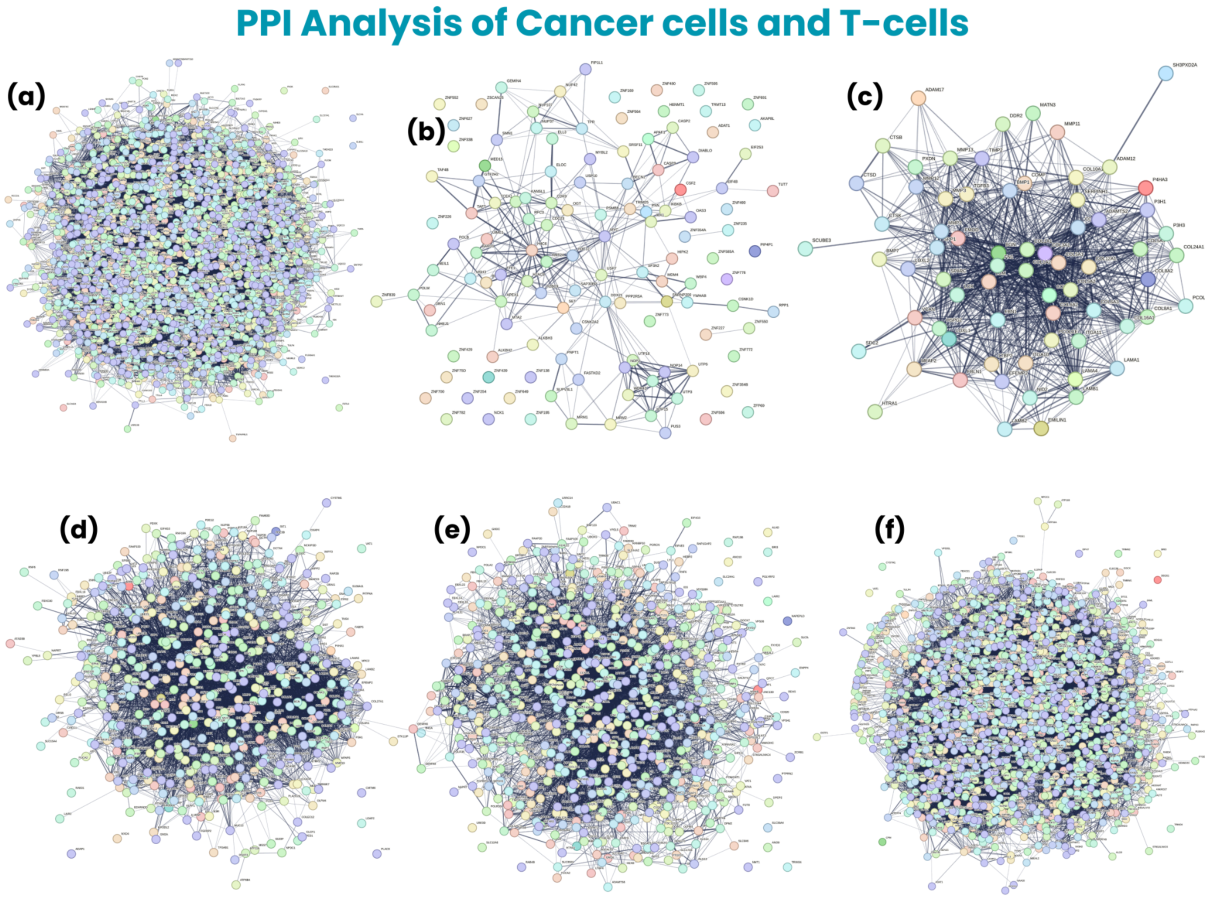


**Figure S5.** The protein-protein interaction analysis showing PPI networks of selected pathway genes from cancer cells and T-cells. **(a)** The PPI network of pathway genes of "common cancer cells genes" group, **(b)** The PPI network of pathway genes of "cancer cells_vs_all-PDAC" group, **(c)** The PPI network of pathway genes of "cancer-PDAC_vs_all-normal" group, **(d)** The PPI network of pathway genes of CD8+ NKT-like cells, **(e)** The PPI network of pathway genes of memory CD4+ T cells, **(f)** The PPI network of pathway genes of naive CD4+ T-cells

**Table S1. The top 10 hub genes of common cancer cell hub genes among "cancer cells_vs_all-PDAC" and "cancer-PDAC_vs_all-normal" groups and unique cancer cell hub genes in both groups along with their number of interactions**

| Common cancer cell hub genes among both groups | |
|---|---|
| **Hub Genes** | **Number of Interactions** |
| GAPDH | 364 |
| AKT1 | 284 |
| EGFR | 242 |
| CS | 212 |
| RHOA | 207 |
| TPI1 | 200 |
| SDHA | 189 |
| TFRC | 189 |
| FASN | 186 |
| HIF1A | 179 |
| **Unique cancer cell genes of "cancer cells_vs_all-PDAC" group** | |
| **Hub Genes** | **Number of Interactions** |
| H4C6 | 29 |
| MYC | 28 |
| H3C12 | 18 |
| DDX21 | 16 |
| USP7 | 16 |
| RFC4 | 12 |
| APEX1 | 12 |
| CDK9 | 12 |
| H2BC9 | 12 |
| NOP2 | 12 |

| Unique cancer cell genes of "cancer-PDAC_vs_all-normal" group | |
|---|---|
| **Hub Gene** | **Number of Interactions** |
| FN1 | 62 |
| COL1A1 | 59 |
| COL1A2 | 56 |
| COL3A1 | 55 |
| COL5A2 | 51 |
| COL6A1 | 50 |
| COL5A1 | 50 |
| BGN | 48 |
| COL6A2 | 46 |
| FBN1 | 45 |

**Table S2. The top 10 hub genes of CD8+ NKT-like cells, memory CD4+ T cells, and naive CD4+ T cells along with their number of interactions**

| **CD8+ NKT-like Cells Hub Genes** | |
|---|---|
| **Hub Gene** | **Number of Interactions** |
| TP53 | 176 |
| FN1 | 170 |
| MMP9 | 149 |
| CD4 | 147 |
| IFNG | 147 |
| NFKB1 | 142 |
| HIF1A | 129 |
| HSP90AA1 | 126 |
| ITGB1 | 119 |
| HSP90AB1 | 116 |
| **Memory CD4+ T Cells Hub Genes** | |
| **Hub Gene** | **Number of Interactions** |
| AKT1 | 200 |
| TP53 | 192 |
| ACTB | 180 |
| CD4 | 148 |
| JUN | 146 |
| FN1 | 142 |
| MMP9 | 126 |
| HSP90AA1 | 123 |
| MAPK3 | 121 |
| HSP90AB1 | 113 |
| **Naive CD4+ T Cells Hub Genes** | |

| **Hub Gene** | **Number of Interactions** |
|---|---|
| TP53 | 452 |
| UBC | 298 |
| UBB | 279 |
| HSP90AA1 | 270 |
| JUN | 260 |
| CTNNB1 | 260 |
| NFKB1 | 251 |
| HSP90AB1 | 238 |
| HSPA8 | 229 |
| H3-3B | 212 |

#### Expression Plots of Cancer cells Hub Genes

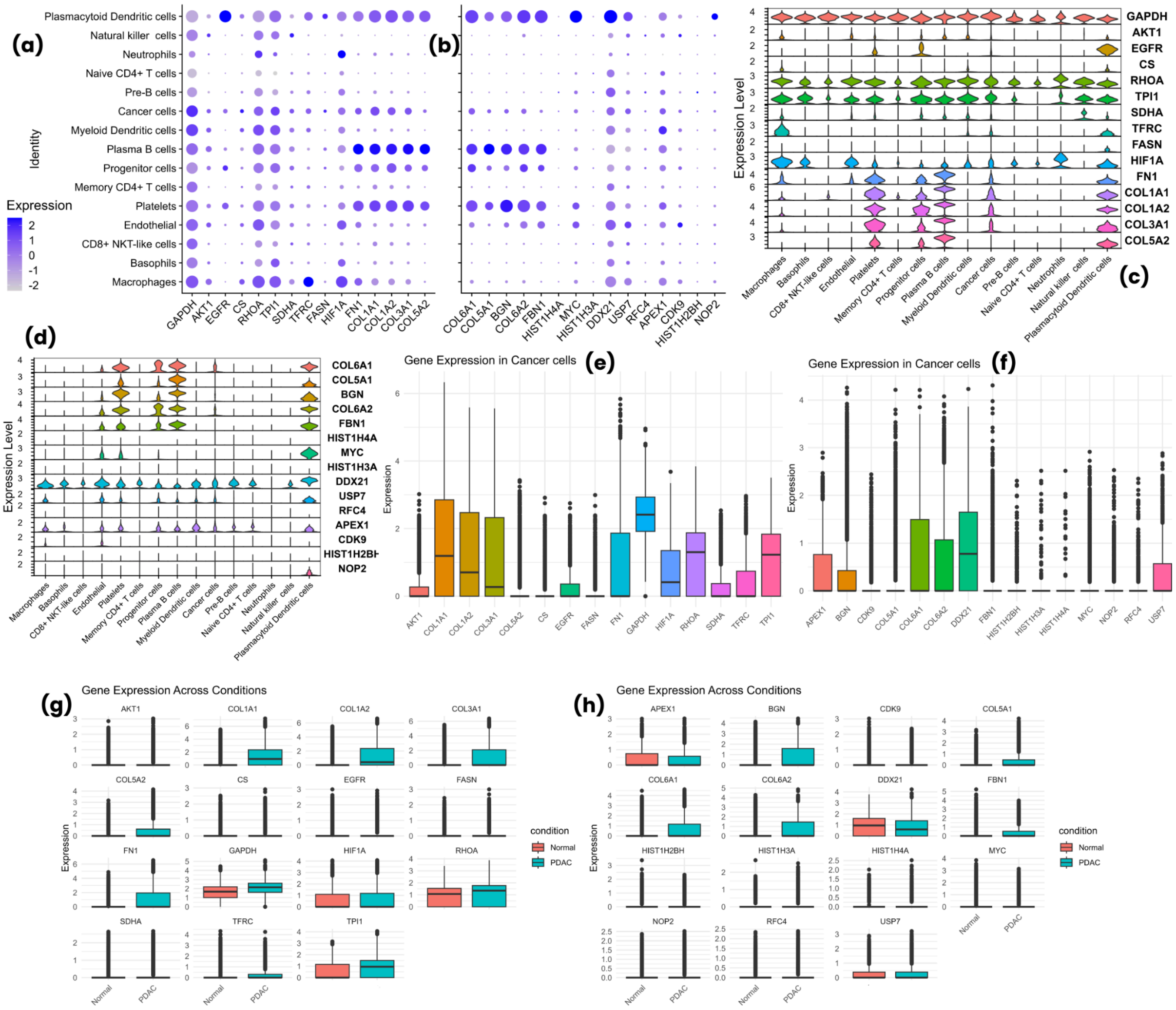


**Figure S6.** Expression levels of all cancer cells groups hub-genes shown in dot plots, violin plots, and box plots, **(a-d)** dot plots and violin plots of cancer cells hub genes showing expression levels in different cell types, **(e-f)** box plots of cancer cell hub genes showing expression levels in cancer cells, **(g-h)** box plots of cancer cell hub genes showing expression levels in conditions

### Expression Plots of CD8+ NKT-like cells and Memory CD4+ T-cells Hub Genes

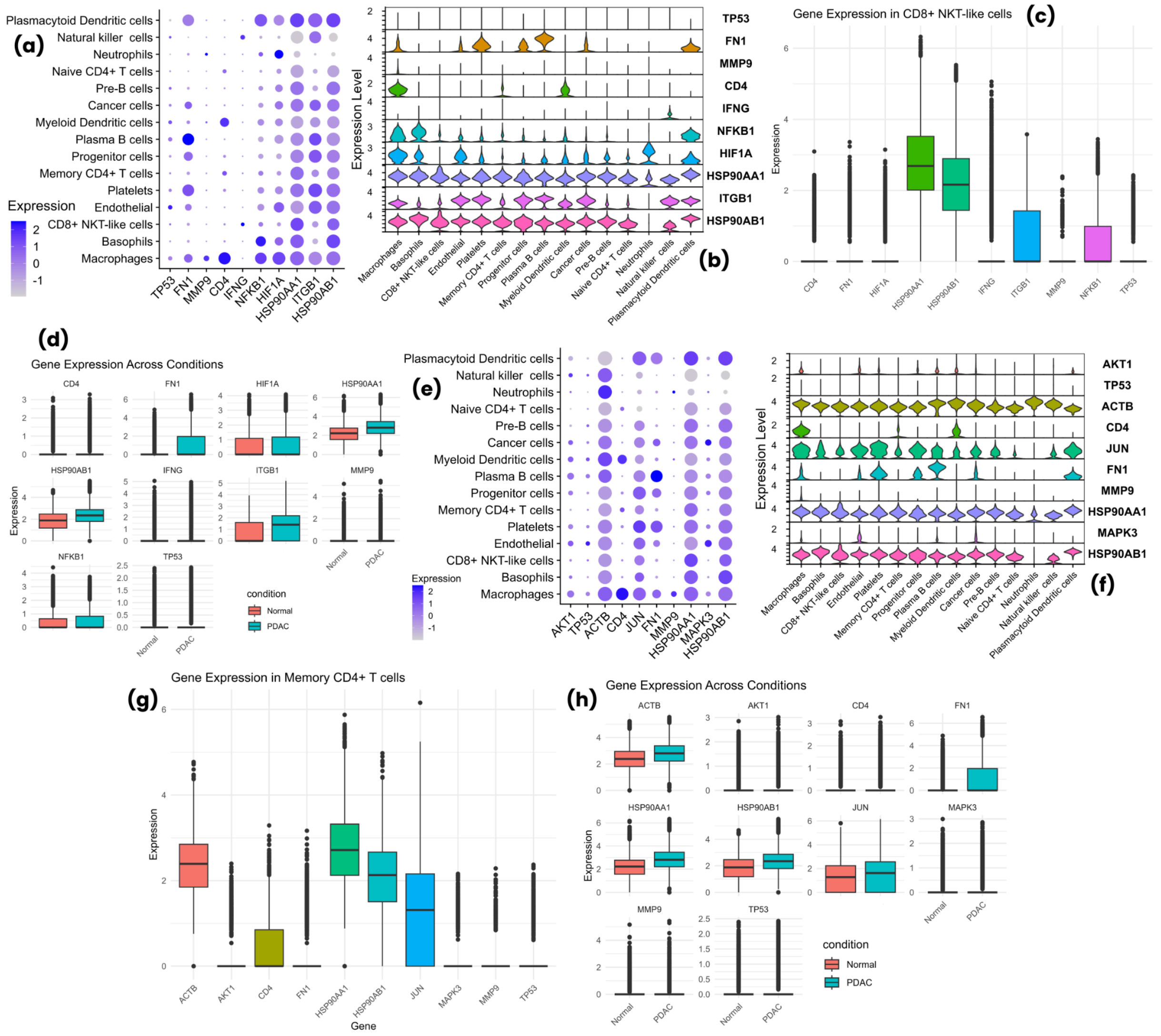


**Figure S7.** Expression levels of T-cells hub genes **(a-d)** Expression levels of CD8+ NKT-like cells hub genes shown in dot plot, violin plot, and box plots across conditions, **(e-h)** expression levels of memory CD4+ T-cells hub genes shown in dot plot, violin plot, and box plots across conditions

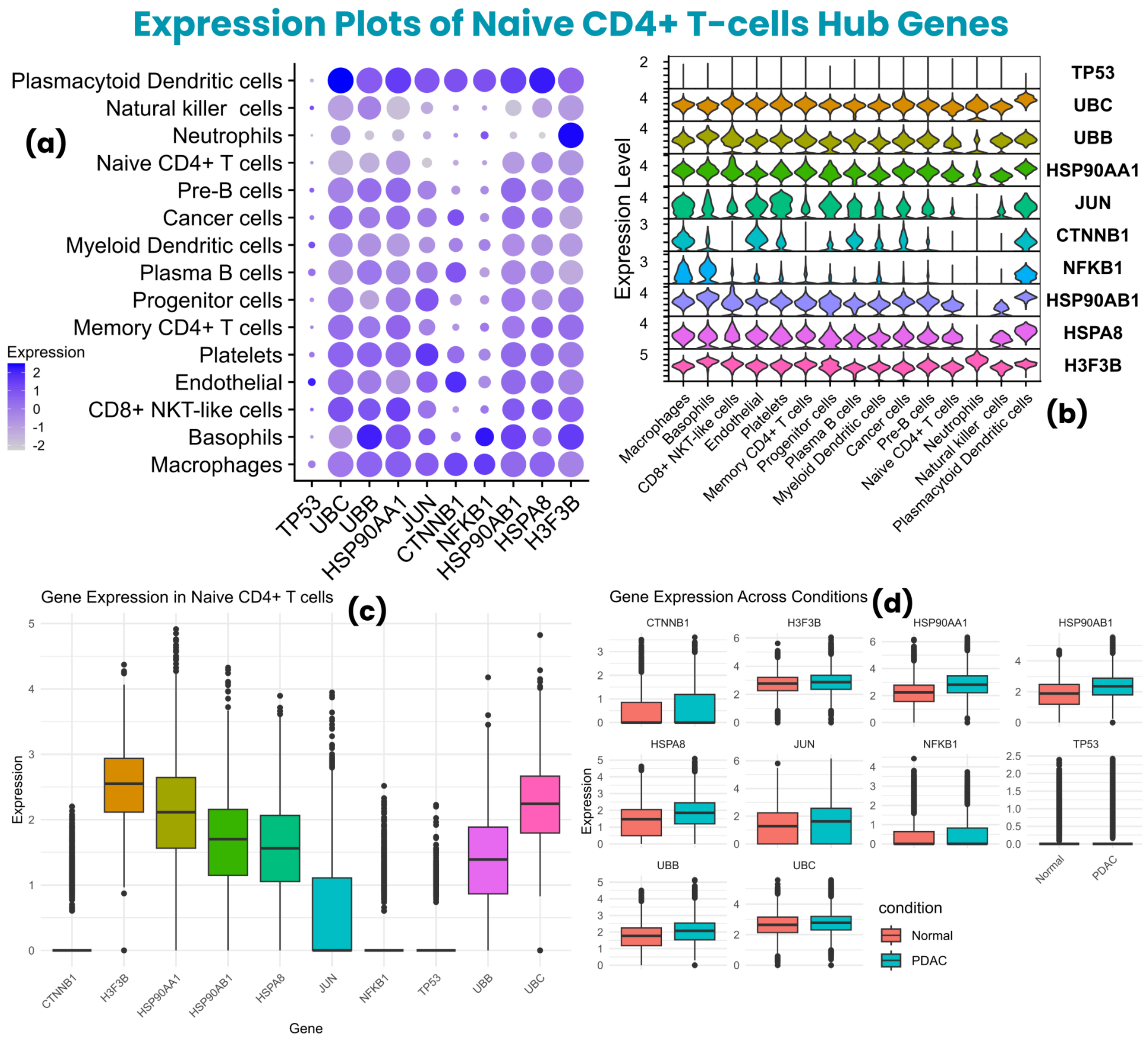


**Figure S8.** Expression levels of T-cells hub genes **(a-d)** Expression levels of naive CD4+ T-cells shown in dot plot, violin plot, and box plots across conditions

**Table S3. Expression levels of cancer cell hub-genes in tumor and control samples mentioned in transcripts per million (TPM) using GEPIA2**

| Common Cancer Cell Hub-Genes | Tumor (TPM) | Normal (TPM) |
|---|---|---|
| GAPDH | 2426.56 | 232.4 |
| AKT1 | 80.5 | 34.08 |
| EGFR | 9.41 | 5.02 |
| CS | 58.68 | 34.33 |
| RHOA | 358.32 | 73.54 |
| TPI1 | 461.79 | 92.66 |
| SDHA | 72.19 | 25.05 |
| TFRC | 24.72 | 7.09 |
| FASN | 43.01 | 13.89 |
| HIF1A | 65.46 | 9 |
| **cancer cells_vs_all-PDAC Hub-Genes** | **Tumor (TPM)** | **Normal (TPM)** |
| H4C6 | 0.17 | 0 |
| MYC | 39.26 | 25.79 |
| H3C12 | 0.07 | 0 |
| DDX21 | 24.08 | 10.96 |
| USP7 | 34.27 | 18.08 |
| RFC4 | 13.18 | 5.83 |
| APEX1 | 105.45 | 43.57 |
| CDK9 | 38.76 | 21.68 |
| H2BC9 | 0.35 | 0 |
| NOP2 | 21.52 | 11.05 |
| **cancer-PDAC_vs_all-normal Hub-Genes** | **Tumor (TPM)** | **Normal (TPM)** |
| FN1 | 978.95 | 17.23 |
| COL1A1 | 1201.65 | 12.61 |
| COL1A2 | 1166.36 | 18.25 |
| COL3A1 | 1103.06 | 13.04 |
| COL5A2 | 71.61 | 1.71 |
| COL6A1 | 179.95 | 35.78 |
| COL5A1 | 115.94 | 2.59 |
| BGN | 613.07 | 22.19 |
| COL6A2 | 277.8 | 30.05 |
| FBN1 | 56.58 | 2.7 |

### Overall Survival Rate of Cancer cells Hub-Genes

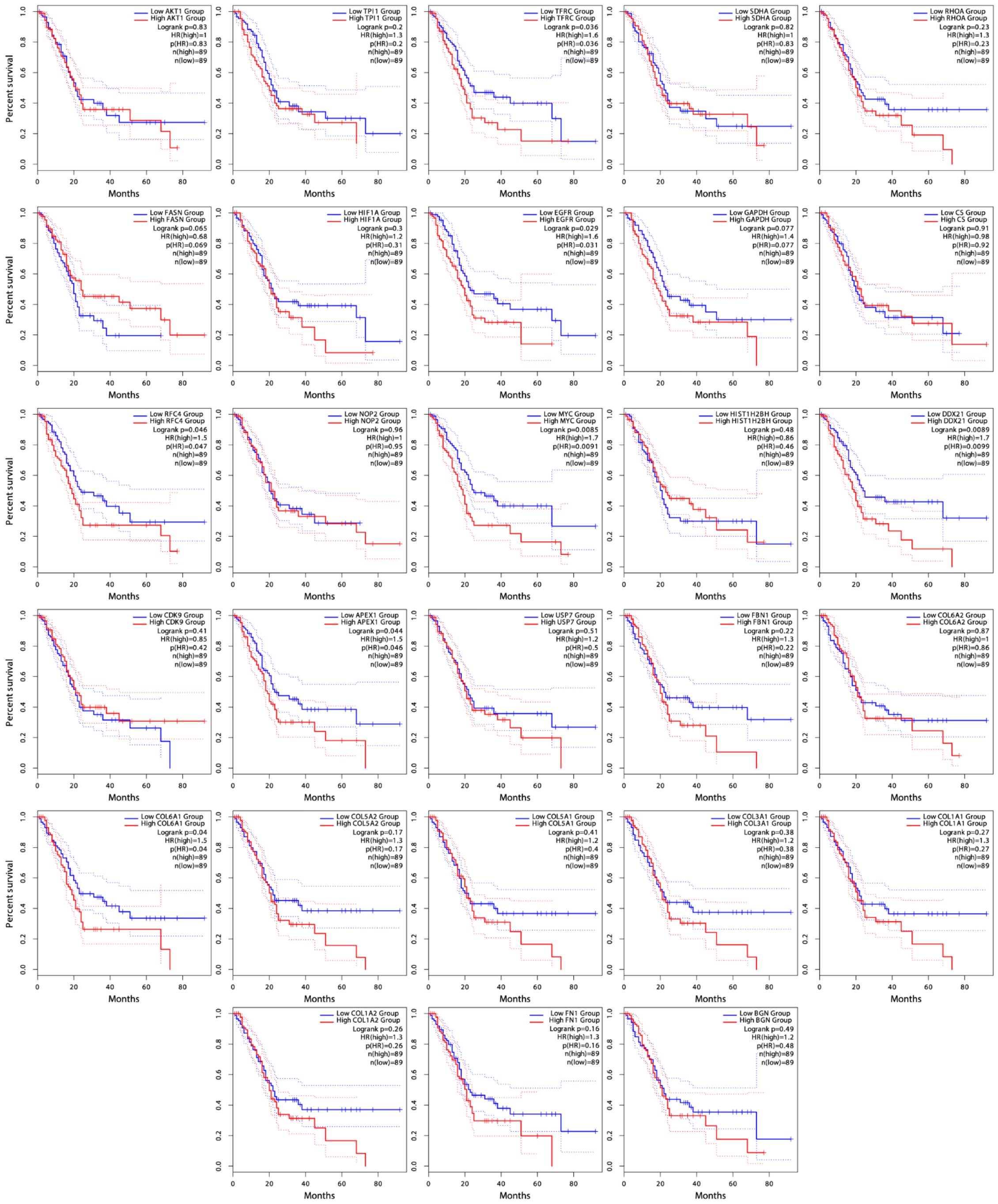


**Figure S9.** The overall survival rate of all significant and insignificant cancer cells hub-genes, indicating the high and low expression of respective hub-gene, leading to overall survival rate in months

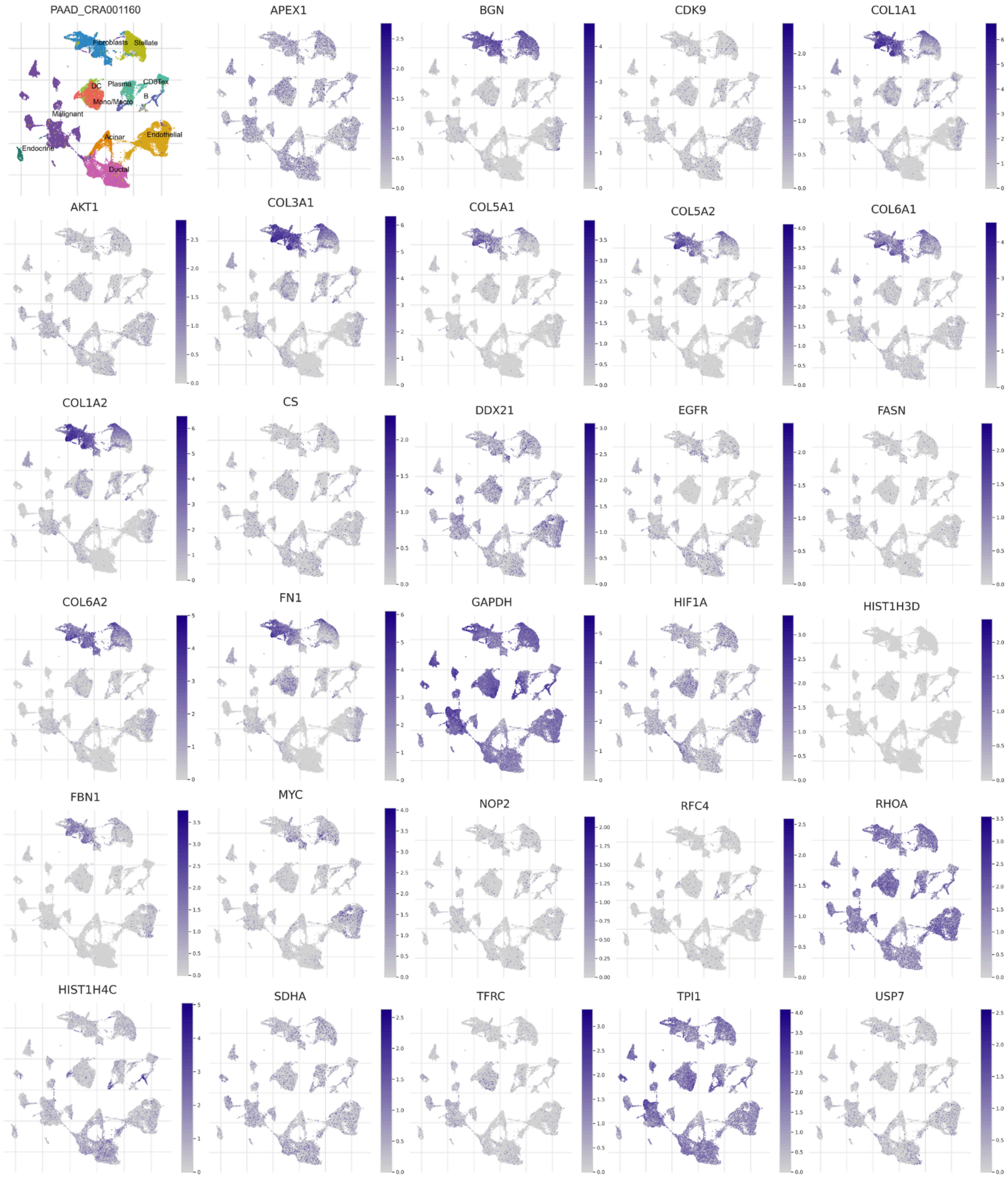


**Figure S10.** Feature plots of each group of cancer cells hub-genes, showing expression levels of each gene in different cell types, including malignant cells

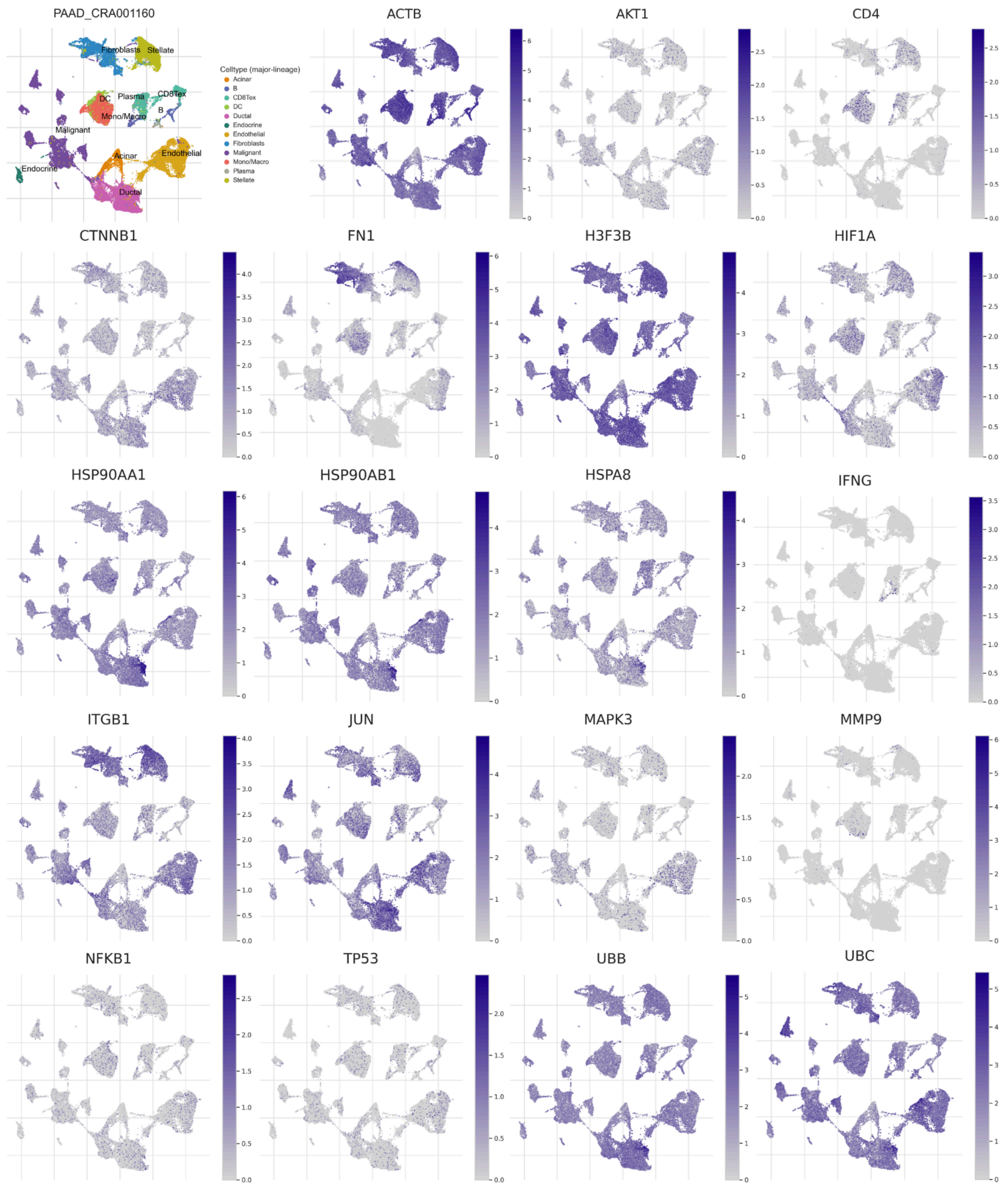


**Figure S11.** Feature plots of T-cells hub-genes showing the expression levels of each gene in different cell types, including CD8Tex cells

#### Expression of Cancer and T-cells Hub-Genes in TISCH2

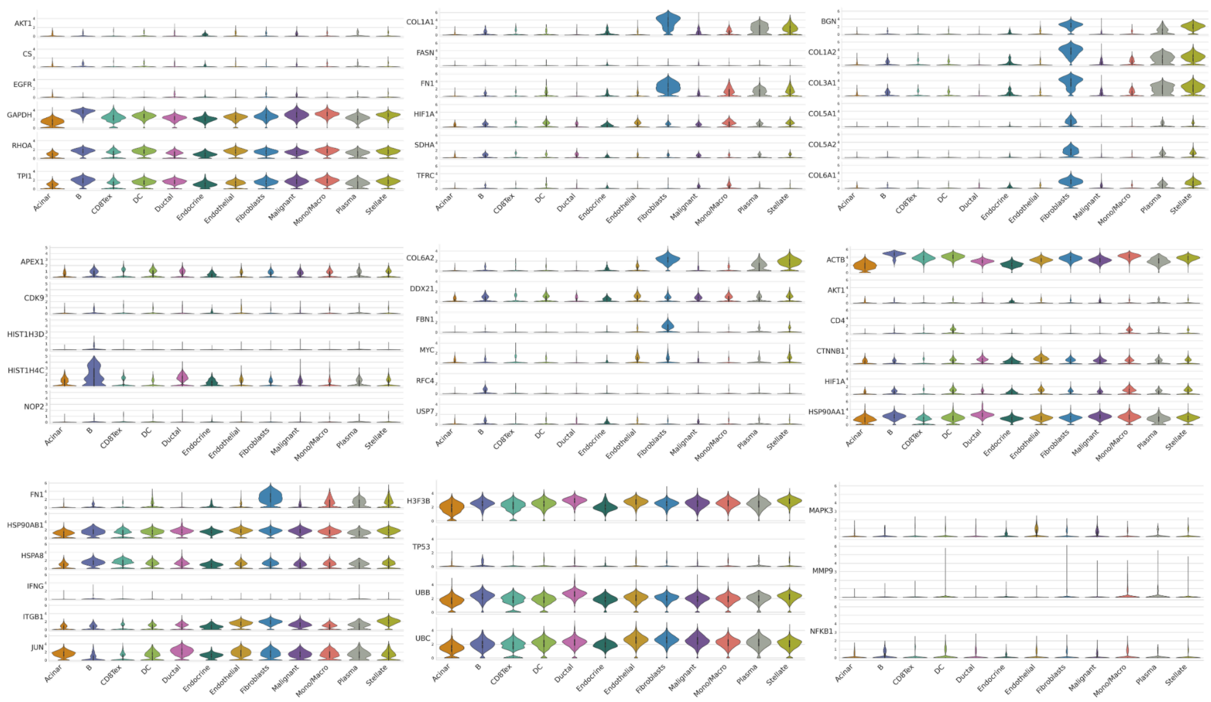


**Figure S12.** Violin plots of cancer and T-cells hub-genes showing expression levels in malignant and CD8Tex cells, along with other cell types within PDAC TIME
