## Supplementary Document 02 for "Novel Insights into T-Cell Exhaustion and Cancer Biomarkers in PDAC Using ScRNA-Seq"

**Single cell RNA sequencing analysis script:**

```
library(dplyr)
library(Seurat)
library(patchwork)
library(openxlsx)
library(HGNChelper)

#Merge samples
sample1 <- Read10X(data.dir = "samples/PDAC/GSM6567157")
sample1 <- CreateSeuratObject(counts = sample1, project = "T1")

sample7 <- Read10X(data.dir = "samples/Normal/GSM6567165")
sample7 <- CreateSeuratObject(counts = sample7, project = "N1")

#Merge different condition samples separately
pdac_combined <- merge(sample1, y = c(sample2, sample3, sample4, sample5, sample6), add.cell.ids = c("T1", "T2", "T3", "T4", "T5", "T6"))

normal_combined <- merge(sample7, y = c(sample8, sample9, sample10, sample11, sample12), add.cell.ids = c("N1", "N2", "N3", "N4", "N5", "N6"))

# Add condition column in the meta.data
pdac_combined <- AddMetaData(object = pdac_combined, metadata = "PDAC", col.name = "condition")

normal_combined <- AddMetaData(object = normal_combined, metadata = "Normal", col.name = "condition")

#Now merge samples with different conditions
combined_samples <- merge(pdac_combined, normal_combined)
dim(combined_samples)

#Load merged data
rawdata <- combined_samples
# The [[ operator can add columns to object metadata. This is a great place to stash QC stats
rawdata[["percent.mt"]] <- PercentageFeatureSet(rawdata, pattern = "^MT-")

# Visualize QC metrics as a violin plot
Plot1 <- VlnPlot(rawdata, features = c("nFeature_RNA", "nCount_RNA", "percent.mt"), ncol = 3)
```

```
#Data Filtration
filtered_data <- subset(rawdata, subset = nFeature_RNA > 200 &
                     nFeature_RNA < 6000 & nCount_RNA > 1000  & percent.mt < 10)
dim(filtered_data)
filter_plt <- VlnPlot(filtered_data, features = c("nFeature_RNA", "nCount_RNA",
"percent.mt"), ncol = 3)

#Data Normalization
normalized_data <- NormalizeData(filtered_data, normalization.method = "LogNormalize",
scale.factor = 10000)

#Highly Variable Features
normalized_data <- FindVariableFeatures(normalized_data, selection.method = "vst",
nfeatures = 3000)

# plot variable features with and without labels
plt1 <- VariableFeaturePlot(normalized_data)
plt2 <- LabelPoints(plot = plt1, points = top10, repel = TRUE)
variable_plt <- plt2

#Data Scaling
scaled_data <- ScaleData(normalized_data)

#Linear Dimensional Reduction
scaled_data <- RunPCA(scaled_data, npcs=50)

# Examine and visualize PCA results a few different ways
print(scaled_data[["pca"]], dims = 1:5, nfeatures = 5)

PCA <- DimPlot(scaled_data, reduction = "pca")

#Determine the Dimensionality of Data
elbow <- ElbowPlot(scaled_data)

#Data Clustering
data <- FindNeighbors(scaled_data, dims = 1:15)
data <- FindClusters(data, resolution = 0.6)

#Non-Linear Dimensional Reduction
data <- RunUMAP(data, dims = 1:15)
data <- RunTSNE(data, dims = 1:15)

#umap
```

```
umap_condition <- DimPlot(data, reduction = "umap", label = TRUE,
                    group.by = "condition")

#TSNE
tSNE_condition <- DimPlot(data, reduction = "tsne",
                    group.by = "condition")

#Cell Annotation (sctype)
# load gene set preparation function
source("https://raw.githubusercontent.com/IanevskiAleksandr/sc-type/master/R/gene_sets_pr
epare.R")
# load cell type annotation function
source("https://raw.githubusercontent.com/IanevskiAleksandr/sc-type/master/R/sctype_score
_.R")

# DB file
db_                                                                                    <-
"https://raw.githubusercontent.com/IanevskiAleksandr/sc-type/master/ScTypeDB_full.xlsx";
tissue <- "Immune system"

# prepare gene sets
gs_list <- gene_sets_prepare(db_, tissue)

# check Seurat object version (scRNA-seq matrix extracted differently in Seurat v4/v5)
seurat_package_v5 <- isFALSE('counts' %in% names(attributes(data[["RNA"]])));
print(sprintf("Seurat object %s is used", ifelse(seurat_package_v5, "v5", "v4")))

# extract scaled scRNA-seq matrix
scRNAseqData_scaled <- if (seurat_package_v5) as.matrix(data[["RNA"]]$scale.data) else
as.matrix(data[["RNA"]]@scale.data)

# run ScType
es.max <- sctype_score(scRNAseqData = scRNAseqData_scaled, scaled = TRUE, gs =
gs_list$gs_positive, gs2 = gs_list$gs_negative)

# merge by cluster
cL_resutls <- do.call("rbind", lapply(unique(data@meta.data$seurat_clusters), function(cl){
                    es.max.cl             =             sort(rowSums(es.max[
,rownames(data@meta.data[data@meta.data$seurat_clusters==cl, ])]), decreasing = !0)
   head(data.frame(cluster = cl, type = names(es.max.cl), scores = es.max.cl, ncells =
sum(data@meta.data$seurat_clusters==cl)), 10)
}))
sctype_scores <- cL_resutls %>% group_by(cluster) %>% top_n(n = 1, wt = scores)
```

```
# set low-confident (low ScType score) clusters to "unknown"
sctype_scores$type[as.numeric(as.character(sctype_scores$scores))< sctype_scores$ncells/4]
<- "Unknown"
print(sctype_scores[,1:3])
write.csv(sctype_scores, file = "Celltypes.csv")

#Overlay the identified cell types on UMAP plot
data@meta.data$sctype_classification = ""
for(j in unique(sctype_scores$cluster)){
  cl_type = sctype_scores[sctype_scores$cluster==j,];
        data@meta.data$sctype_classification[data@meta.data$seurat_clusters == j] =
as.character(cl_type$type[1])
}

#Visualize annotated data
#umap and tsne
umap_dis <- DimPlot(data, reduction = "umap", label = TRUE, repel = TRUE, group.by =
'sctype_classification', split.by = 'condition')

tsne_dis <- DimPlot(data, reduction = "tsne", label = TRUE, repel = TRUE, group.by =
'sctype_classification', split.by = 'condition')

##### Differential between conditions ####
data$celltype.condition <- paste(data$sctype_classification, data$condition, sep = "_")
Idents(data) <- "celltype.condition"

Naive_CD4_markers <- FindMarkers(data, ident.1 = "Naive CD4+ T cells_PDAC",
                                ident.2 = "Naive CD4+ T cells_Normal", verbose = TRUE, only.pos =
FALSE)

Naive_CD4_Up_markers <- subset(Naive_CD4_markers, avg_log2FC > 0.25)
Naive_CD4_Dn_markers <- subset(Naive_CD4_markers, avg_log2FC < -0.25)

#Find markers between cell types in same condition
Cancer_markers <- FindMarkers(data_joined, ident.1 = "Cancer cells",
                        ident.2 = NULL, test.use = "wilcox",
                        only.pos = FALSE)

Cancer_Up <- subset(Cancer_markers, avg_log2FC > 0.25)
Cancer_down <- subset(Cancer_markers, avg_log2FC < -0.25)
```

**Molecular subtypes classification script for PDAC tumor samples:**

```
# UCell Analysis
signatures <- list(
  classical = c("ELF3", "NFIX", "CUX1", "SSBP3", "GATA4", "GATA6", "MUC1",
              "FOXA1", "FOXA2", "FOXA3", "HES1", "HNF4A", "HNF4G",
              "HNF1A", "HNF1B", "MUC5AC", "LAMA1", "PDX1", "MNX1",
              "BMP2", "FOS", "FOXP1", "FOXP4", "KLF4", "SHH"),

  basal_like = c("IL1A", "CXCL1", "VEGFA", "SEMA3C", "S100A2", "KRT6A", "LY6D",
               "LDHA", "TP63", "ZEB1", "ZEB2", "TWIST", "GLI1", "GLI2",
               "SNAI1", "SNAI2", "SLC16A3", "TPI1", "ENO1", "HIF1A",
               "FOXM1", "NSDHL", "CD7", "EGFR", "TP53", "MYC", "MYBL1"),

    immunogenic = c("IDO1", "ADORA2A", "BTLA", "CD27", "CD40LG", "CD48",
"CTLA4",
                "ICOS", "ICOSLG", "IDO2", "LAG3", "LAIR1", "PDCD1", "PDCD1LG2",
                "TIGIT", "TNFRSF25", "TNFRSF4", "TNFRSF8", "VSIR", "CD80",
                "CD86", "HAVCR2", "TNFRSF18", "TNFRSF9", "CD28", "CD40",
                "CALR", "IFNB1", "MET", "EIF2AK4", "P2RY2", "LRP1", "EIF2A",
                "ANXA1", "PANX1", "PLG", "COPS5", "FYN", "IRF3", "ITGB3",
                "SPTA1", "CD3D", "CD3E", "CD4", "CD8", "CD25", "FOXP3",
                "PD-L1", "TLR4", "TLR7", "TLR8", "PD-L2", "CSF1R", "PD1"),

  adex = c("HOXA3", "CDH3", "LIMK1", "SLC17A7", "SLC25A5", "SLC35A2", "RBPJL",
         "INS", "NEUROD1", "MAFA", "NKX2-2", "MIST1", "NR5A2", "AMY2B",
         "PRSS1", "PRSS3", "CEL", "ELA3A", "CFTR"),

   normal_activated_stroma = c("ACTA2", "VIM", "DES", "ITGAM", "CCL13", "CCL18",
"PDGFRB",  "FAP", "SPARC", "WNT2", "WNT5A"))

data_ucell <- AddModuleScore_UCell(data_subset, features = signatures, name = "_UCell",
                                   ncores = 2)

vln_plt <- VlnPlot(data_ucell, features = signatures, group.by = "orig.ident")


feat_plt <- FeaturePlot(data_ucell, reduction = "tsne",
                        features = signatures, ncol = 3, order = T)
```
